# *Pantoea agglomerans* T6SS effectors reside in hotspots with combinatorial offensive and defensive arsenals

**DOI:** 10.64898/2026.08.01.741415

**Authors:** Ksenia Leo, Chaya Mushka Fridman, Anushree Haldar, Tridib Mahata, Udi Qimron, Eran Bosis, Dor Salomon

## Abstract

Gram-negative bacteria deploy type VI secretion systems (T6SSs) to mediate interbacterial competition. Although numerous T6SS effectors have been identified, their pan-genomic repertoires and evolutionary dynamics remain poorly understood. Here, we combine proteomics and comparative genomics to map the T6SS effector landscape across *Pantoea agglomerans*, a diverse species that includes pathogenic and beneficial strains. We uncover an extensive pan-genomic arsenal in which most effectors are encoded outside the main T6SS gene cluster, within highly dynamic hotspots distributed across the chromosome and megaplasmids. Analysis of these hotspots reveals multilayered, combinatorial arrangements of shuffled genetic cargo. Remarkably, these loci act as versatile “genomic armories” that co-localize offensive antibacterial weapons with protective anti-phage defense systems. By investigating uncharacterized genes within these variable regions, we discovered and validated a novel T6SS effector and a previously unknown anti-phage defense system, named Juno. Collectively, our findings demonstrate that bacterial warfare arsenals are highly modular and reside in dynamic genomic hubs that alternate or combine interbacterial aggression with viral defense. This evolutionary association reveals a functional blurring between offensive and defensive strategies within the bacterial accessory genome. Our findings further highlight orphan effector-associated variable regions as promising leads in the search for unrecognized bacterial conflict and defense systems.

## INTRODUCTION

Many Gram-negative bacteria employ the type VI secretion system (T6SS), a protein delivery apparatus, to outcompete rivals (1). T6SSs deploy toxic effectors into neighboring cells in a contact-dependent manner. The effectors decorate a tube-spike complex comprising a tube of stacked Hcp hexamers capped by a VgrG trimer and a PAAR repeat-containing protein that sharpens it (2). This complex is propelled out of the cell by the contraction of an engulfing sheath (3).

Effectors are categorized as either “specialized”, comprising a toxic domain fused to one of the secreted tube-spike components (i.e, Hcp, VgrG, or PAAR), or “cargo”, including toxic proteins that are loaded onto the secreted tube-spike non-covalently (4), often via an adapter (5–7) or co-effector (8). Although several T6SSs were shown to target eukaryotes (9–15), most T6SSs investigated to date target bacteria by delivering a cocktail of antibacterial effectors that manipulate conserved bacterial cell components, such as nucleic acids (e.g., DNases (16–19)), energy balance (e.g., NADases (20, 21) and (p)ppApp synthases (22)), membranes (e.g., phospholipases (23) and pore-forming toxins (24–27)), peptidoglycan (e.g., amidases (28), glycoside hydrolases (29), and l, d-transpeptidases named Tlde (30)), cell division machinery (31), protein translation (32, 33), and DNA replication (34). Notably, antibacterial effectors are encoded next to a cognate immunity protein that antagonizes their toxicity, thereby preventing self- and kin-intoxication (4).

Many effectors reside in evolutionary dynamic islands within clusters that encode all or most of the structural components of the T6SS apparatus (35–38). Effectors may also reside in auxiliary operons, often as part of mobile genetic elements (MGEs) (16, 39–42), partially accounting for the plasticity of T6SS effector repertoires; in this work, we will refer to these as orphan effectors (6, 43).

T6SS effector repertoires differ between bacterial species and even between closely related strains of the same species (24, 27, 35, 40, 42, 44). In several bacteria, the effector repertoire was described as comprising: (i) core effectors, found in all or nearly all examined genomes of the species containing a T6SS of interest; and (ii) accessory effectors, found only in a subset of genomes containing a T6SS of interest (9, 42, 45). However, whether the presence of core effectors is a characteristic shared among all T6SSs remains unexplored. Moreover, it is unclear whether orphan effectors are distributed randomly across the genome or are concentrated in hotspots.

Although the T6SS and its effector repertoire have been thoroughly characterized in many animal pathogens, these systems remain poorly understood in phytopathogens (46–49). *Pantoea agglomerans*, a Gram-negative bacterium within the family Enterobacteriaceae, is ubiquitously found across diverse natural and agricultural habitats (50, 51). This species exhibits a diverse ecological profile: while some strains are phytopathogens that target various plant species, others serve as beneficial biocontrol agents, and certain isolates have recently emerged as opportunistic human pathogens (52).

In previous work, we investigated T6SS1 in *Pantoea agglomerans* pv. *betae* (*Pab*), a phytopathogen that induces tumor-like galls in beet and gypsophila (53–55). We found that T6SS1, homologs of which are present in many *Pantoea* and *Erwinia* species (36, 56), mediates interbacterial competition against various Gram-negative bacteria under standard laboratory conditions (35). Furthermore, we described three effector-harboring genetic islands within the T6SS1 main gene cluster that differ among *Pantoea* strains (35) and identified two orphan effectors (named Pse3 and Pse4) with N-terminal PIX delivery domains through computational analyses (43). Thus, *Pab* is known to deliver at least six effectors: the specialized VgrG and PAAR-Rhs effectors, and the cargo effectors Pse1, Pse2, Pse3, and Pse4 (35, 43).

In the current work, we set out to determine whether *Pab* encodes additional orphan T6SS1 effectors and, if so, to characterize their genomic locations and conservation within the species. Using comparative proteomics and computational analyses, we identify 17 additional orphan effectors in the *Pab* genome. We show that although none of the 21 antibacterial cargo effectors encoded by *Pab* is conserved across all *P. agglomerans* strains that harbor a similar T6SS, three effector types are represented in ∼90% of strains. Furthermore, we demonstrate that all orphan T6SS effectors reside in variable genomic regions, some of which constitute genomic hotspots on the main chromosome and megaplasmids. These hotspots harbor diverse armories comprising defensive and offensive tools, either together or interchangeably.

## RESULTS

### Pab harbors an extensive T6SS1 effector arsenal

Aiming to identify the T6SS1 effector repertoire of *Pab*, we performed a comparative proteomics analysis (9, 45, 57). Using mass spectrometry, we compared the proteins secreted by the wild-type *Pab* (T6SS1^+^) with those secreted by a Δ*tssA* mutant strain (T6SS1^−^ (35)) when grown in 2xYT media at 28°C. We identified 19 proteins significantly enriched in the supernatant of the wild-type strain (Fig. 1A and Dataset S1). These include the known tube-spike secreted structural components Hcp, VgrG, and PAAR (the latter two are specialized effectors (35)), and the previously reported cargo effectors Pse1 and Pse2, encoded within the main T6SS1 gene cluster (35). The other 14 identified proteins are predicted T6SS cargo effectors; they were named *<u>P</u>antoea* type <u>s</u>ix <u>e</u>ffector 6-14 (Pse6-Pse14), as explained below, in accordance with previous nomenclature (35, 43) (Table 1).

**Figure 1.**
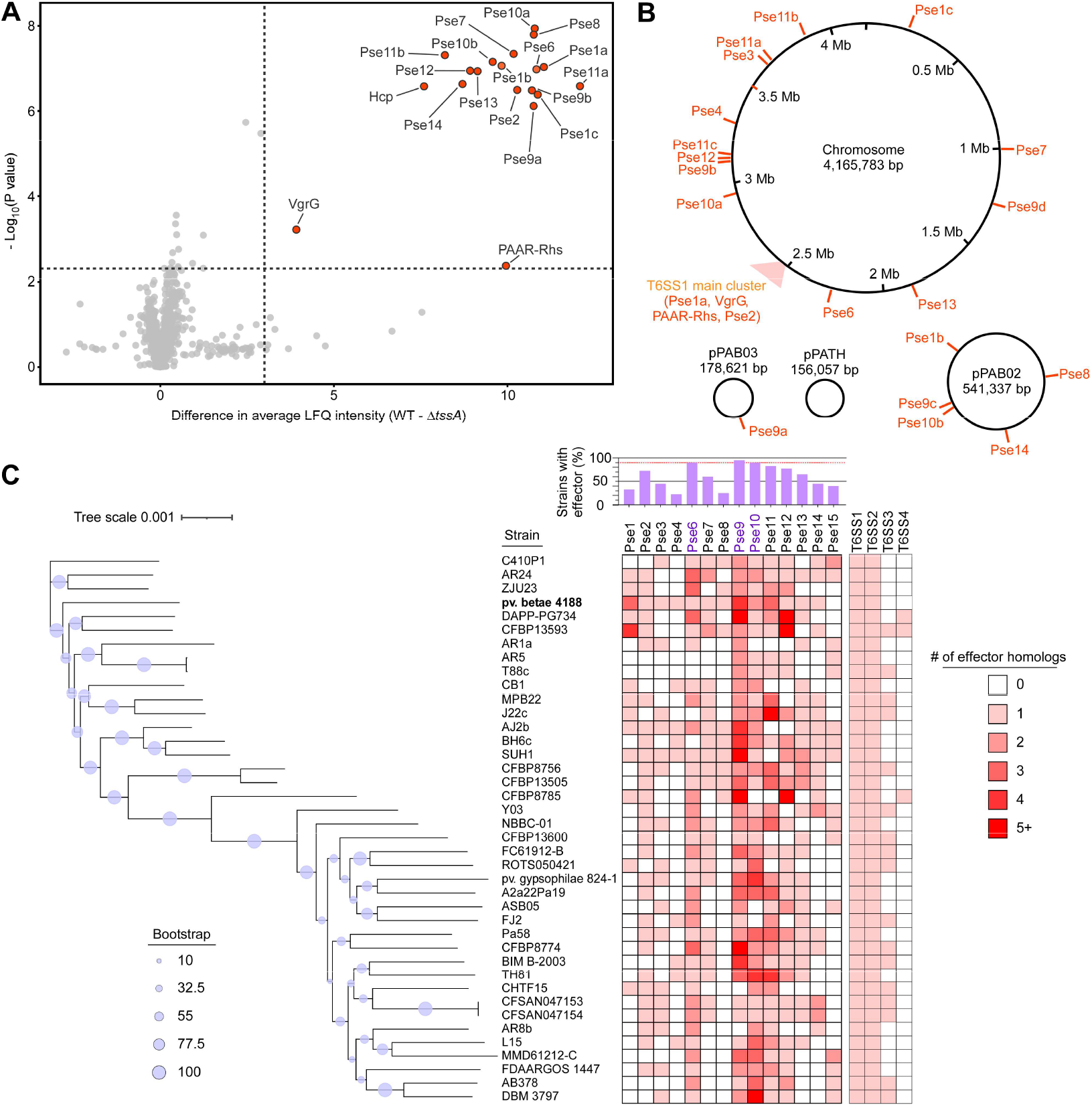
*Pab* harbors orphan T6SS effectors that are not conserved in other *P. agglomerans* strains. **(A)** Volcano plot summarizing the comparative analysis of proteins identified in the media of wild-type (WT) *Pab* and a T6SS1^−^ mutant (Δ*tssA*) strain, using label-free quantification (LFQ). The average LFQ signal intensity difference between the WT and Δ*tssA* strains is plotted against the −Log10 of Student’s *t*-test *P*-values (*n* = 3 biological replicates). Proteins that were significantly more abundant in the secretome of the WT strain (difference in average LFQ intensities > 3; *P*-value < 0.005; at least one unique peptide in each biological replicate) are denoted in red. **(B)** Schematic representation of effectors’ location within the *Pab* genome. **(C)** Heatmaps of effector type and T6SS cluster distribution in 40 complete *P. agglomerans* genomes. The phylogenetic tree is based on a comparison of 1, 000 concatenated core proteins across the indicated strains. The evolutionary history was inferred using the maximum likelihood method. Bootstrap values appear on the corresponding branch as percentages across 1, 000 replicates.

**Table 1.**
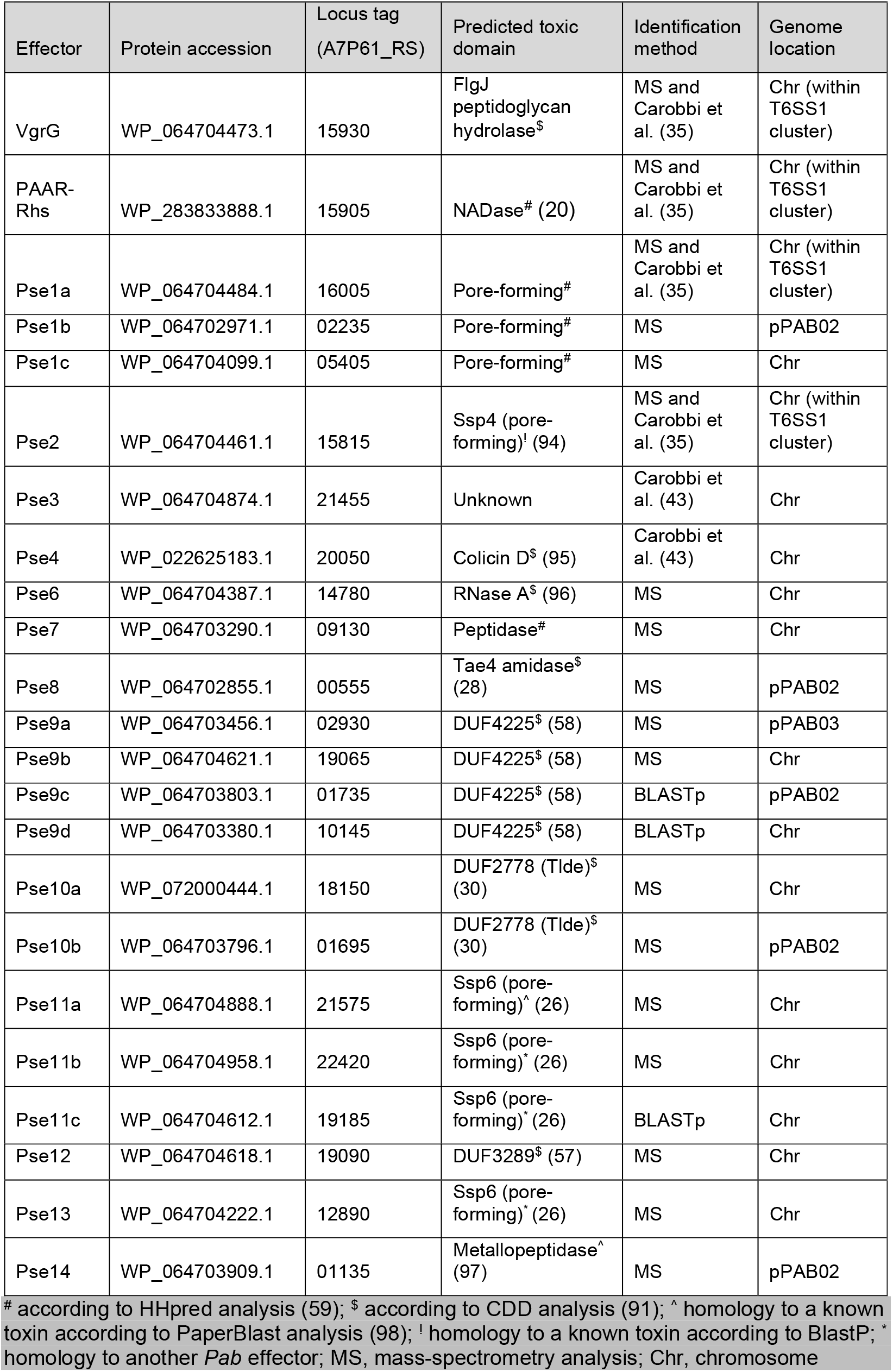
T6SS effectors in Pab.

When analyzing the effectors identified using the proteomics approach, we noticed that several are homologous. For example, WP_064702971.1 and WP_064704099.1 are homologs of the cluster-encoded Pse1 (35). Therefore, we named them Pse1b and Pse1c, respectively, and renamed the cluster-encoded effector Pse1a. Other homologous effectors contain a known toxic domain previously associated with T6SS effectors, e.g., Pse9a and Pse9b, containing DUF4225 (58); Pse10a and Pse10b, containing a DUF2778 domain, also known as Tlde (30); and Pse11a and Pse11b, which are homologs of the pore-forming effector Ssp6 from *Neisseria* (26) (Table 1). The presence of homologous effectors within the same genome led us to investigate whether additional homologs not detected by the proteomics analysis are present in *Pab*. Indeed, we identified three genes encoding homologs of the identified effectors: two additional DUF4225-containing proteins (Pse9c and Pse9d) and one similar to Ssp6 (Pse11c) (Table 1).

All of the newly identified effectors, aside from Pse7, are either homologs of previously described antibacterial effectors from other bacteria (e.g., Pse11, Pse13, and Pse14) or contain a predicted domain that had previously been assigned an antibacterial toxic activity (e.g., Pse6, Pse8, Pse9, and Pse10), suggesting that they are antibacterial T6SS effectors (Table 1). Notably, we did not detect non-T6SS secretion signals, such as Sec or Tat signal peptides, among the identified effectors. Moreover, all but one (Pse11b) have a neighboring small gene encoding a known or predicted immunity protein (*<u>P</u>antoea* type <u>s</u>ix Immunity, Psi), and some homologs of these predicted immunity proteins are also encoded in multiple copies (e.g., two *psi3* in tandem) or elsewhere in the genome (e.g., a Psi9 homolog encoded downstream of *psi7*).

### Pse7 is a bona fide antibacterial T6SS1 effector

Because Pse7 is not similar to any known secreted antibacterial toxin, we sought to validate its identity as an antibacterial T6SS effector. Hidden Markov model analysis suggested that Pse7 contains a C-terminal peptidase domain of the M1 family, with a conserved HExxH motif (84% probability according to HHpred (59); amino acids 165-258 of Pse7 compared to amino acids 24-105 of PF01433 Pfam domain). To the best of our knowledge, M1 peptidases have not been identified as T6SS effectors. To confirm that Pse7 is secreted in a T6SS1-dependent manner, we monitored the secretion of a plasmid-encoded, C-terminal Myc-tagged Pse7 from a wild-type strain and a T6SS1-inactive mutant strain (Δ*tssA*). Consistent with the proteomics results, Pse7 was secreted into the growth medium only by the wild-type strain (Fig. S1A). To determine whether it is a bona fide antibacterial T6SS1 effector and whether the downstream-encoded protein Psi7 is its cognate immunity, we deleted *pse7* and *psi7* and used the mutant strain (Δ*psei7*) as prey in self-competition assays. We reasoned that if Pse7 and Psi7 are a T6SS effector and immunity pair, their deletion would render the prey strain sensitive to attack by a wild-type *Pab* strain that delivers the effector via T6SS1, and not by a T6SS1^−^ mutant strain. Indeed, we found that deleting this gene pair renders the prey strain sensitive to a T6SS1-mediated attack, as evidenced by the T6SS-dependent decrease in prey viability after 5 hours of co-incubation (Fig. S1B). Importantly, the expression of Psi7 from a plasmid restored the sensitive prey strain’s ability to antagonize the T6SS1-mediated attack. Taken together, these results confirm that Pse7 and Psi7 constitute a T6SS1 antibacterial effector-immunity pair.

In light of the predicted activities of the effectors and the confirmed role of *Pab* T6SS1, we conclude that the newly identified and predicted effectors are antibacterial. When considering previously identified effectors not observed in the current comparative proteomics analysis (i.e., Pse3 and Pse4 (43)), *Pab* T6SS1 carries the largest effector arsenal described to date, comprising at least 23 effectors.

### T6SS1 orphan effector distribution in P. agglomerans genomes

While four of the identified *Pab* T6SS1 effectors are located within the effector islands of the main T6SS1 cluster (i.e., Pse1a, Pse2, VgrG, and PAAR-Rhs) (35), the rest are located outside the main cluster, and thus, we consider them orphan effectors. Of these, 13 are encoded on the chromosome, five on the pPAB02 megaplasmid, and one on the pPAB03 plasmid (Fig. 1B). No T6SS effectors are found on the pPATH plasmid carrying the virulence-associated type III secretion system (60). Notably, Pse9d is encoded next to *hcp* (WP_064703379.1; 67% amino acid sequence identity with the Hcp encoded within the main T6SS1 gene cluster), as previously observed for other DUF4225-containing effectors (58).

To determine the pan-genome distribution of T6SS1 cargo effectors identified in *Pab*, we analyzed their occurrence by effector type (e.g., the homologs Pse1a, Pse1b, and Pse1c were collectively analyzed as Pse1) across 40 publicly available, complete RefSeq genomes confirmed by OrthoANI analysis as *P. agglomerans* (Dataset S2 and Dataset S3). Notably, all 40 genomes harbor the conserved T6SS1 and T6SS2 clusters (T6SS2 is a partial yet conserved cluster of unknown function (56)), whereas the distribution of the previously described T6SS3 and an additional cluster, which we named T6SS4, is patchy (36, 56) (Fig. 1C, Fig. S2, and Dataset S3). Interestingly, no effector type was identified across all *P. agglomerans* genomes. Nevertheless, three effector types (i.e., Pse6, Pse9, and Pse10) are present in at least 90% of the investigated genomes; thus, we consider them the core cargo effector repertoire of T6SS1 (42, 45). Other effectors are less frequent and display a patchy distribution (Fig. 1C). Notably, many genomes harbor multiple homologs of the same effector type (e.g., up to 11 homologs of Pse12 in *P. agglomerans* DAPP-PG734) (Dataset S3).

### Pab T6SS effectors reside within genomic hotspots

Because none of the orphan *Pab* T6SS1 effectors are present in all 40 complete genomes, we hypothesized that they reside in variable genomic regions that harbor diverse cargoes. To examine this, we compared sequence conservation between the *Pab* genome and the 39 other complete *P. agglomerans* genomes described above (Fig. 2). Alignment of the 100-kb regions flanking each orphan *Pab* effector revealed that all effectors reside within regions of low sequence conservation (Fig. 3). These findings suggest that orphan *Pab* T6SS effectors are replaced by alternative sequences in other *P. agglomerans* genomes and therefore reside within variable genomic regions (VRs) (61–64).

**Figure 2.**
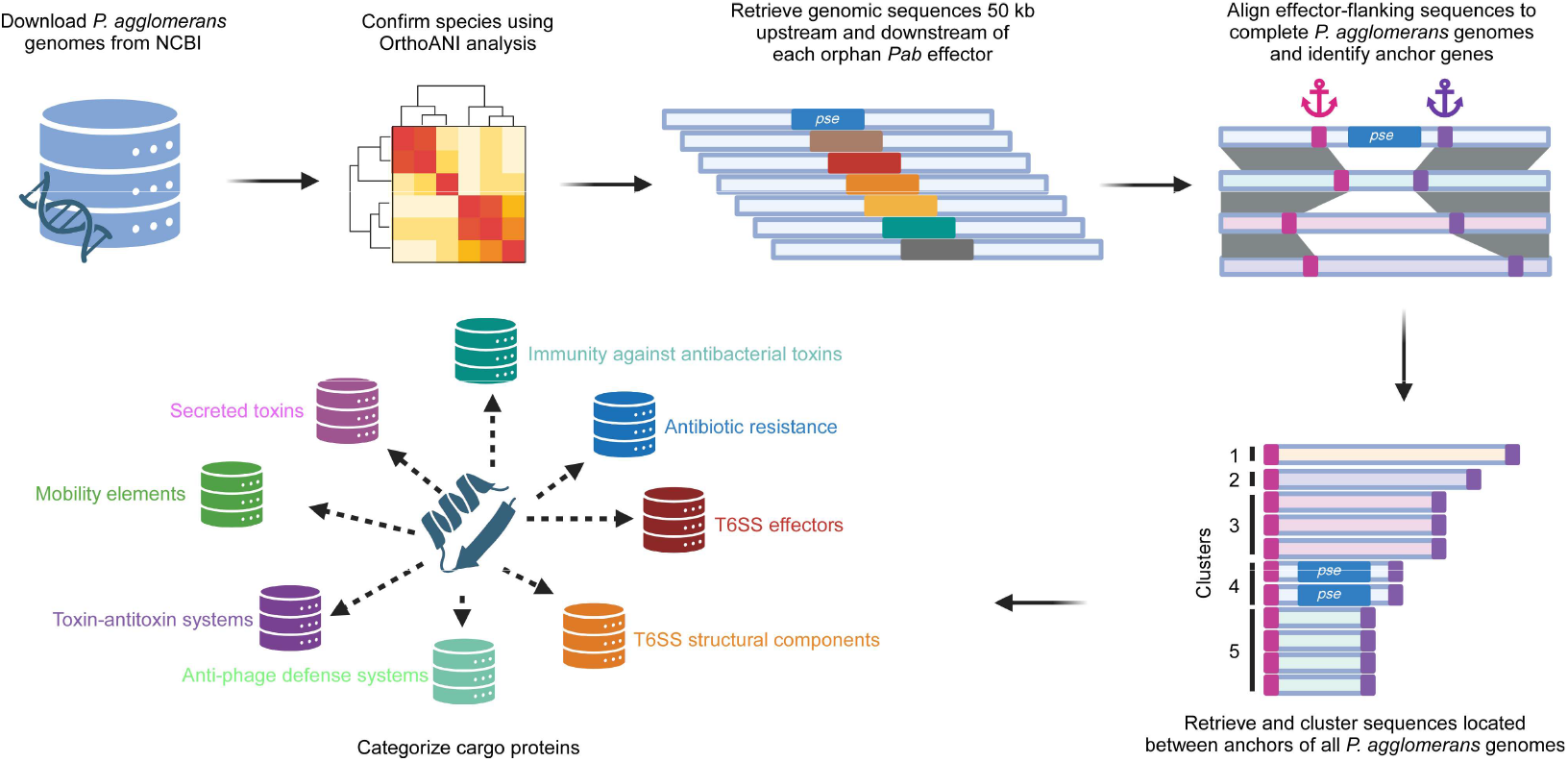
Variable genomic region analysis workflow. A schematic representation of the workflow used to identify and analyze the cargo within *Pab* variable genomic regions that harbor T6SS effectors. The figure was prepared using BioRender.com.

**Figure 3.**
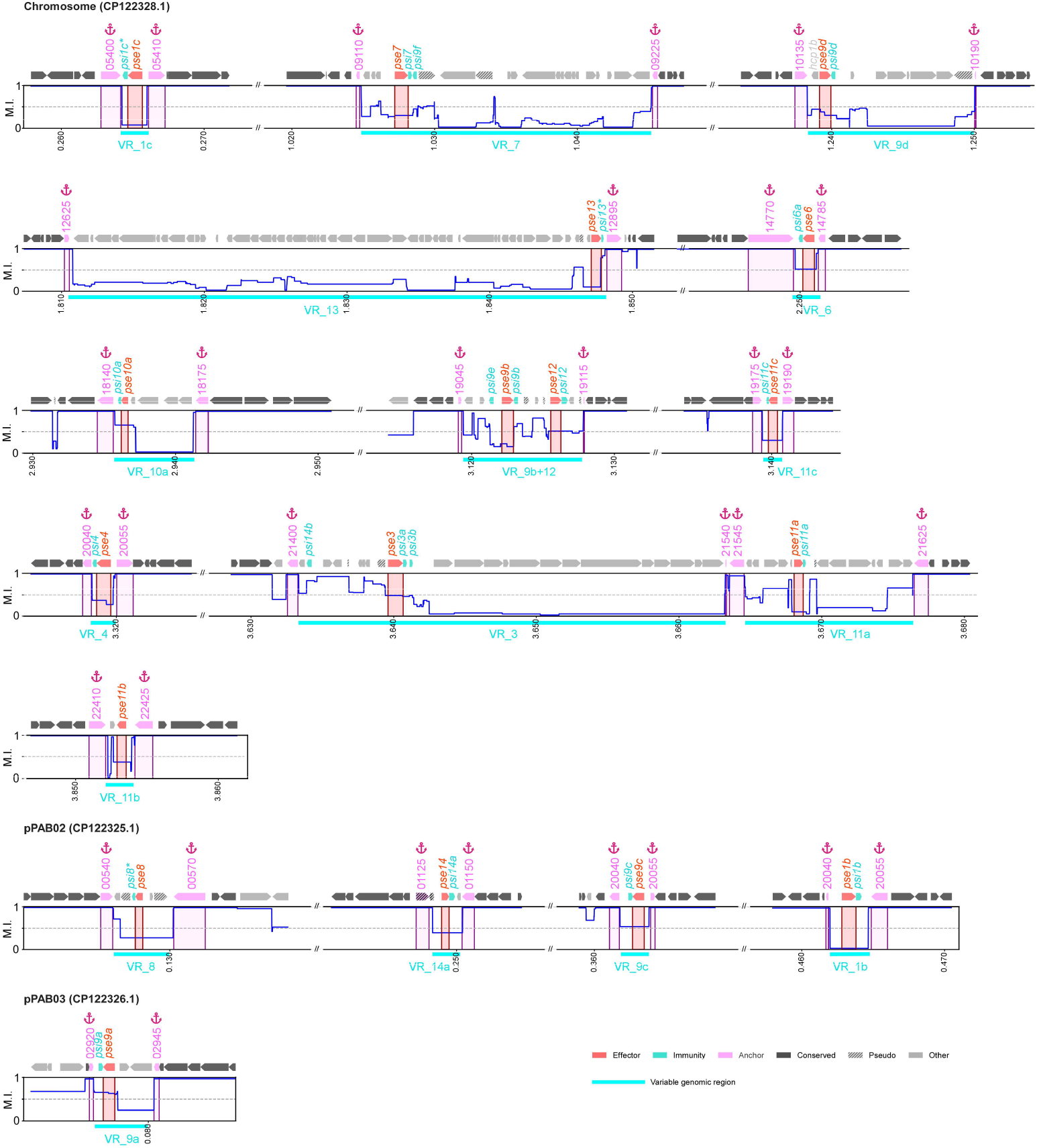
*Pab* T6SS1 effectors reside within regions of low sequence conservation. Sequence conservation analysis across 40 complete *P. agglomerans* genomes, shown as the mean identity (M.I.; blue line) at *Pab* T6SS1 effector-harboring locations. Conserved genes are those present in all 40 complete *P. agglomerans* genomes with a similarity index ≥ 0.8. Locus (A7P61_RSxxxxx) or gene names in *Pab* are shown above; *Pab* genome positions are shown below. Asterisks denote manually annotated immunity genes.

To identify the sequences that replace the *Pab* effectors in other genomes, we first sought to determine the borders of the VRs that contain each *Pab* orphan T6SS effector. To this end, we identified flanking genes conserved in both sequence and relative orientation across the 40 complete *P. agglomerans* genomes (Fig. 2 and Fig. 3; details are available in the Methods section); the first conserved genes upstream and downstream of each effector served as anchors for identifying cargo sequences. We identified anchor genes for 18 of the 19 orphan *Pab* effectors (Fig. 3 and Table 2). However, we were unable to confidently identify conserved flanking genes for *pse10b*, and therefore excluded it from subsequent cargo analyses. We also noticed that *pse9b* and *pse12* reside within the same VR (VR_9b+12), as they share the same set of anchors (Fig. 3 and Table 2).

**Table 2.** Anchors identified for *P. agglomerans* variable genomic regions.

| Variable genomic region | Left anchor locus (A7P61_RS) | Left anchor product | Right anchor locus (A7P61_RS) | Right anchor product |
| --- | --- | --- | --- | --- |
| VR_1b | 02230 | GhoT/OrtT family toxin | 02250 | lactaldehyde reductase |
| VR_1c | 05400 | MFS transporter | 05410 | HD-GYP domain-containing protein |
| VR_3 | 21400 | CDP-diacylglycerol diphosphatase | 21540 | tRNA-Leu |
| VR_4 | 20040 | methylthioribulose 1-phosphate dehydratase | 20055 | pyridoxal phosphate-dependent aminotransferase |
| VR_6 | 14770 | multidrug efflux RND transporter permease subunit OqxB | 14785 | RrF2 family transcriptional regulator |
| VR_7 | 09110 | YmjA family protein | 09225 | transfer-messenger RNA |
| VR_8 | 00540 | VOC family protein | 00570 | ferric-rhodotorulic acid/ferric-coprogen receptor FhuE |
| VR_9a | 02920 | type II toxin-antitoxin system RelE family toxin | 02945 | 4-amino-4-deoxy-L-arabinose-phosphoundecaprenol flippase subunit ArnF |
| VR_9b+12 | 19045 | DUF2171 domain-containing protein | 19115 | tRNA-Arg |
| VR_9c | 01725 | hypothetical protein | 01740 | hypothetical protein |
| VR_9d | 10135 | helix-turn-helix domain-containing protein | 10190 | tRNA-Arg |
| VR_10a | 18140 | suppressor of fused domain protein | 18175 | oxygenase MpaB family protein |
| VR_10b | 01685 | Bor family protein | 01700 | ArsR/SmtB family transcription factor |
| VR_11a | 21545 | DUF1176 domain-containing protein | 21625 | YsnF/AvaK domain-containing protein |
| VR_11b | 22410 | Gfo/Idh/MocA family protein | 22425 | MFS transporter |
| VR_11c | 19175 | multifunctional acyl-CoA thioesterase I/protease I/lysophospholipase L1 | 19190 | SDR family oxidoreductase |
| VR_13 | 12625 | YebY family protein | 12895 | alcohol dehydrogenase AdhP |
| VR_14 | 01125* | LysR family transcriptional regulator | 01150 | metal ABC transporter permease |

\* The left anchor for VR\_14 was manually changed from A7P61\_RS01130 to the adjacent gene, A7P61\_RS01125, because A7P61\_RS01130 is a partial duplication of the right anchor.

To characterize the different cargoes found within the 17 identified VRs in the *P. agglomerans* pan-genome, we searched for the anchor genes of each VR in a database of 363 publicly available RefSeq *P. agglomerans* genome sequences, which were confirmed by OrthoANI analysis (Dataset S2). We then extracted the sequences located between the corresponding anchors in each genome. Clustering of the extracted cargo sequences (using CD-HIT-EST (65); ≥ 85% sequence identity and coverage) revealed that 13 VRs contain cargoes of diverse lengths that could be assigned to 6-160 sequence clusters, indicating that these regions constitute genomic hotspots (Fig. 4). In contrast, only two possible cargoes are found in four VRs (i.e., VR_1b, VR_1c, VR_9c, and VR_14; Fig. 4A). These include either the effector and immunity-containing cargo found in *Pab* (i.e., harboring *Pse1b*, *Pse1c*, *Pse9c*, and *Pse14*) or an alternative sequence lacking annotated genes (Dataset S4).

**Figure 4.**
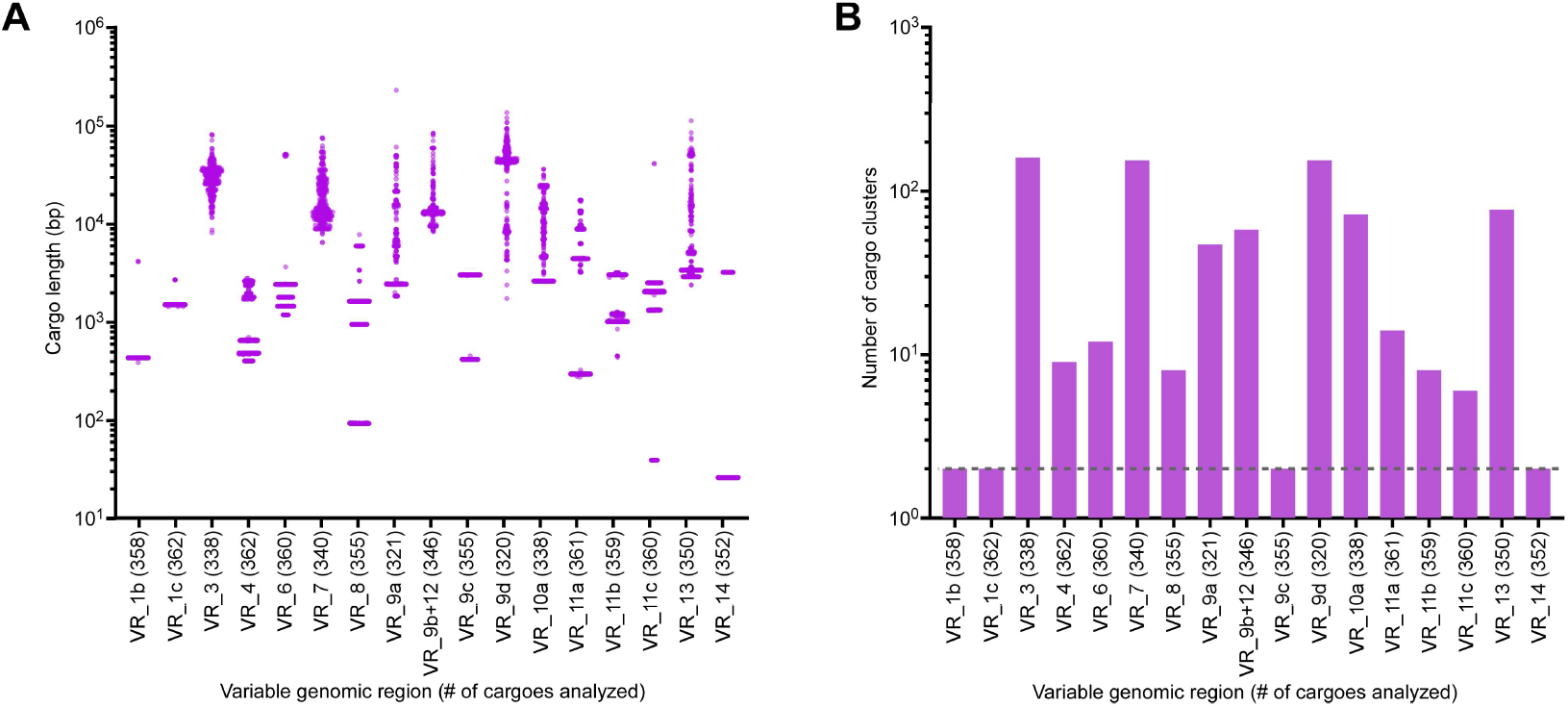
Many *P. agglomerans* variable regions are hotspots containing diverse cargo. **(A)** Length distribution of cargoes extracted from the specified variable genomic regions (VR) in *P. agglomerans* RefSeq genomes. **(B)** The number of cargo sequence clusters found in each VR described in A. A dashed line denotes two sequence clusters.

### P. agglomerans hotspots comprise “genomic armories”

Next, we analyzed the genes within each cargo. Using a combination of in-house and publicly available databases and servers (see Methods section for details), we assigned each gene to a specific category of interest that has previously been identified in bacterial MGEs and affect bacterial fitness: T6SS structural components, T6SS effectors, immunity against antibacterial toxins, secreted toxins, anti-phage defense systems, toxin-antitoxin systems, antibiotic resistance, mobility elements, or others (Fig. 2 and Dataset S4).

While analyzing the T6SS1 effectors within the extracted cargoes, we noticed that none of the VRs harbors a T6SS effector across all *P. agglomerans* genomes (Fig. 5A-B and Dataset S4). This supports our initial observation that orphan *Pab* T6SS effectors are not conserved within the *P. agglomerans* pan-genome (Fig. 1C). Furthermore, we found that several VRs harbor diverse repertoires of T6SS effectors across the *P. agglomerans* pan-genome, and that, occasionally, multiple effectors are present within the same cargo (Fig. 5C, Fig. S3-S5, and Dataset S4). This observation indicates that the arsenal of T6SS effectors within the VRs is dynamic and suggests that effectors are shuffled among them.

**Figure 5.**
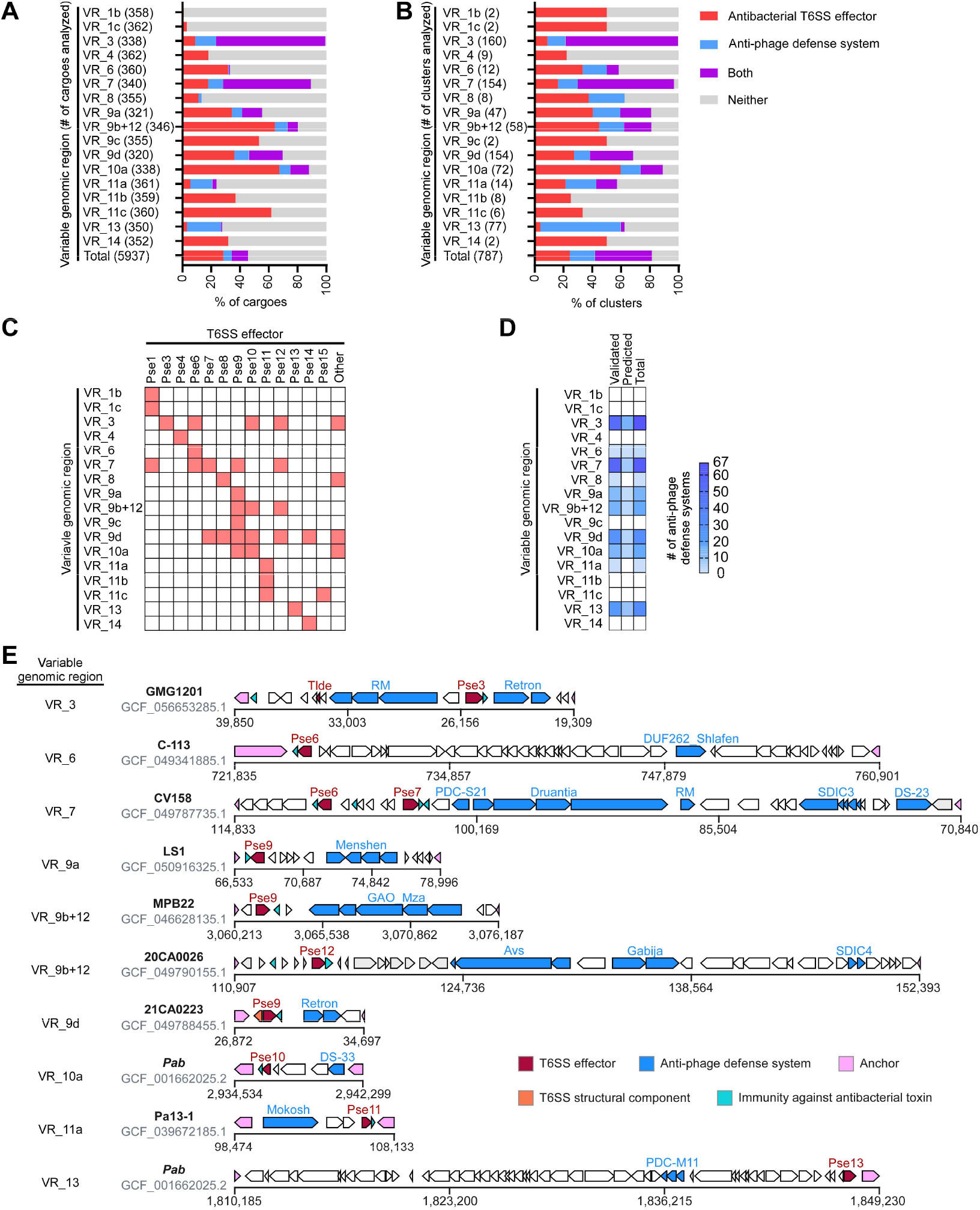
*P. agglomerans* hotspots contain antibacterial T6SS effectors and anti-phage defense systems. **(A-B)** Percentage of *P. agglomerans* variable genomic region cargoes (A) or cargo sequence clusters (B) harboring antibacterial T6SS effectors, anti-phage defense systems, or both. **(C)** Heatmap of the antibacterial T6SS effectors identified in *P. agglomerans* variable genomic regions. **(D)** Heatmap showing the number of distinct anti-phage defense systems, validated or predicted, identified in *P. agglomerans* variable genomic regions. The analyses include the anti-phage defense system Juno and the T6SS effector Pse15, which were identified in this study and are detailed below. **(E)** Representative cargoes of variable genomic regions containing both antibacterial T6SS effectors (red) and anti-phage defense systems (blue). *P. agglomerans* strain names and NCBI RefSeq assembly numbers are denoted on the left.

We also found that 10 of the 13 VRs designated as hotspots (i.e., comprising > 2 sequence clusters) harbor diverse anti-phage defense systems expected to protect the bacteria from invading bacteriophages and other MGEs (66) (Fig. 5A-B, Fig. 5D, Fig. S3-S6, and Dataset S4). Four VRs (i.e., VR_3, VR_7, VR_9d, and VR_13) contain a diverse and dynamic repertoire of > 30 different types of anti-phage defense systems within their cargoes (Fig. 5D); altogether, we identified 118 validated and 37 predicted anti-phage defense systems (according to DefenseFinder (67) and PADLOC (68) databases) within the VRs that we analyzed (Fig. S7), suggesting that these hotspots are hubs for anti-phage defense systems. Remarkably, in 9 of the hotspots, cargoes harbor anti-phage defense systems alongside antibacterial T6SS effectors, rendering these “genomic armories” with combined offensive and defensive arsenals (Fig. 5E, Fig. S3-S6, and Dataset S4).

In addition to antibacterial T6SS effectors and anti-phage defense systems, “genomic armory” VRs in *P. agglomerans* harbor additional mechanisms tailored for inter-microbial conflict. For example, many cargoes include orphan immunity genes (i.e., not adjacent to a cognate antibacterial toxin or effector) that are predicted to antagonize antibacterial toxins delivered into the bacterium (e.g., T6SS effectors or bacteriocins) (Fig. S3-S5 and Dataset S4). Moreover, antibiotic resistance genes are found within the cargoes of VR_9a, VR_9b+12, VR_9d, and VR_10a (Fig. 6A and Dataset S4), thereby providing an additional layer of defense.

**Fig. 6.**
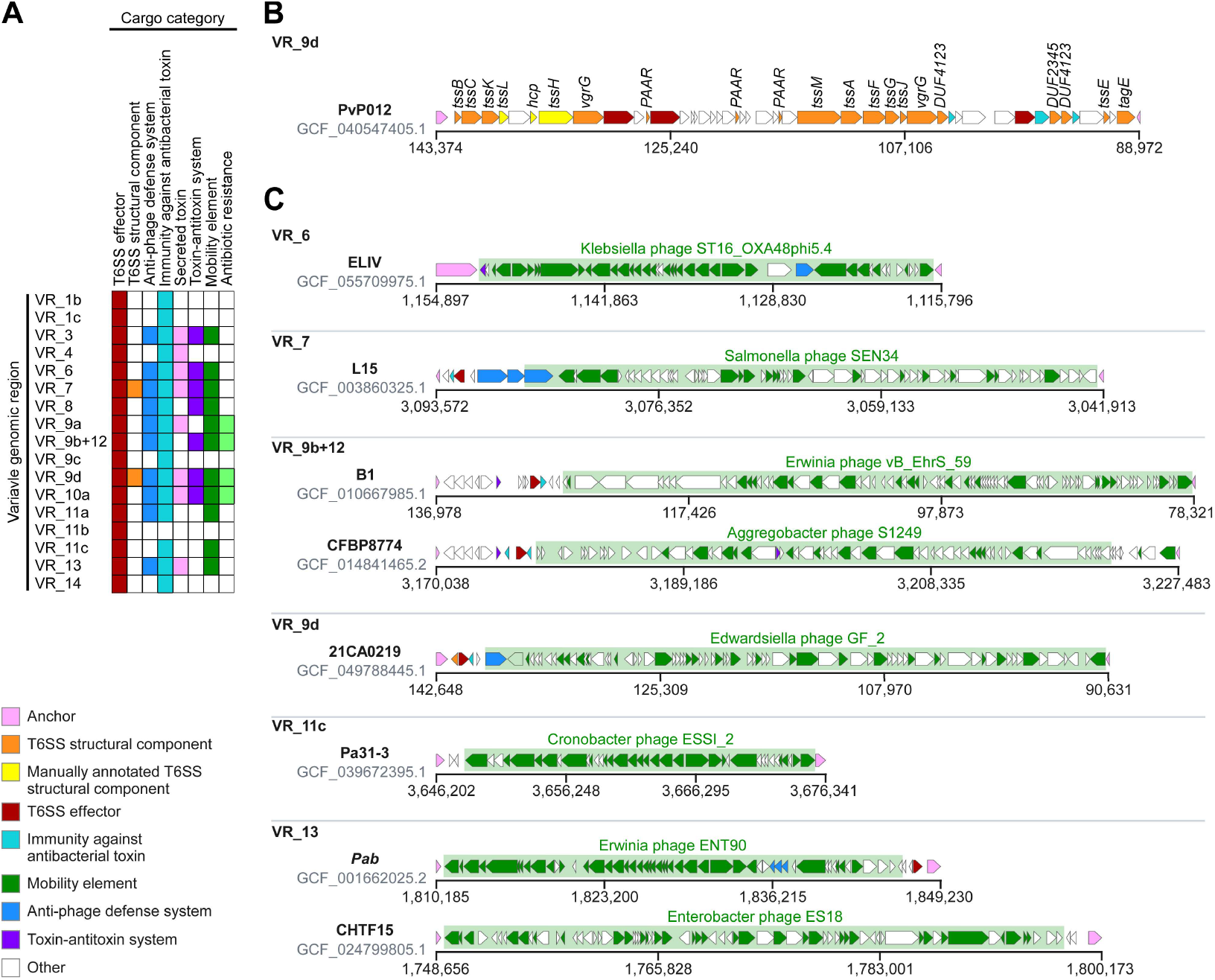
Certain *P. agglomerans* hotspots harbor complete T6SS gene clusters or prophages. **(A)** Heatmap showing the presence of different categories in the cargoes of each variable genomic region. **(B)** A representative VR_9d cargo containing a T6SS3 gene cluster. **(C)** Representative cargoes of variable genomic regions containing prophages. The prophage regions (as predicted by PHASTEST) are denoted by a green rectangle with the predicted bacteriophage name above it. *P. agglomerans* strain names and RefSeq assembly numbers are denoted on the left.

### Effectors can be replaced by complete T6SS clusters and prophages

As described above, the VRs that contain T6SS1 effectors in *Pab* are often replaced by diverse, alternative cargoes in other *P. agglomerans* strains. Interestingly, we found that in some strains, Pse9d, which is encoded within the VR_9d of *Pab*, is replaced by a complete T6SS3 gene cluster that is accompanied by its own repertoire of antibacterial T6SS effectors, including: WHIX effector family members (57), FIX effector family members (16), phospholipases (28), and homologs of *Pab* T6SS1 effectors such as Pse10 and Pse14 (Fig. 6B and Dataset S4).

Several VRs, including VR_6, VR_7, VR_9b+12, VR_9d, VR_11c, and VR_13, harbor a complete prophage (according to PHASTEST analysis (69)) in their cargo (Fig. 6C and Dataset S5). Notably, in VR_9b+12 and VR_13, two different types of prophage cargo were observed (Fig. 6C and Dataset S5). These prophages may serve as additional defensive or offensive mechanisms, as other prophages have previously been shown to mediate defense against invading bacteriophages (70) and to serve as inducible weapons during competition between bacterial populations (71).

Collectively, our observations reveal that *P. agglomerans* VRs serve as “genomic armories” harboring complex layers of offensive and defensive arsenals, including complete T6SS clusters and prophages, individual antibacterial T6SS effectors and anti-phage defense systems, and antibiotic resistance genes. These tools appear together or interchangeably within the different VRs.

### Using P. agglomerans hotspots to identify additional T6SS effectors

Because the *P. agglomerans* VRs are hubs for antibacterial T6SS effectors, we reasoned that additional effectors could be identified by investigating their cargoes. The cargoes within VR_11c of the different *P. agglomerans* genomes cluster into six sequence groups (Fig. 4B). In one of these groups, the cargo comprising *pse11c* and *psi11c* in *Pab* is replaced by a cargo comprising two alternative genes organized in a bicistronic unit resembling a T6SS effector-immunity pair (e.g., in *P. agglomerans* DSM3493T; Fig. S6). The first gene encodes a protein with a predicted C-terminal pesticin domain (WP_062758233.1), which was previously associated with peptidoglycan-hydrolyzing T6SS effectors (57); the second gene encodes a protein with a predicted Lysozyme inhibitor LprI domain (WP_022626094.1), which we previously predicted is used in immunity proteins, since it is often found within the *Pab* T6SS1 cluster islands (35). In light of these predicted activities, we hypothesized that these two genes constitute a T6SS1 effector-immunity pair and renamed them p*se15* and *psi15*, respectively.

Analysis aimed at determining the presence of Pse15 across the 40 complete *P. agglomerans* genomes revealed a patchy distribution in < 50% of the genomes (Fig. 1C). Since a Pse15 homolog was not identified in the *Pab* genome, we set out to employ *Pab* as a surrogate T6SS1 platform and investigate whether exogenous expression of *pse15* and *psi15* will enable this strain to outcompete a parental *Pab* strain lacking these genes during co-incubation (a methodology that we used previously in vibrios (58)). In accordance with our hypothesis, plasmid-borne *pse15* and *psi15* from *P. agglomerans* NBBC-01 enabled *Pab* to outcompete its parental prey strain in a T6SS1-dependent manner, compared to an empty plasmid (Fig. 7A). This is evident from a positive competition index, calculated as the ratio of prey CFUs collected at 5 h after co-incubation with a T6SS^−^ (Δt*ssA*) attacker strain to those at the beginning of the experiment, divided by the same ratio for co-incubation with the wild-type attacker strains. Taken together, our results indicate that Pse15 is another *P. agglomerans* antibacterial effector delivered by T6SS1.

**Figure 7.**
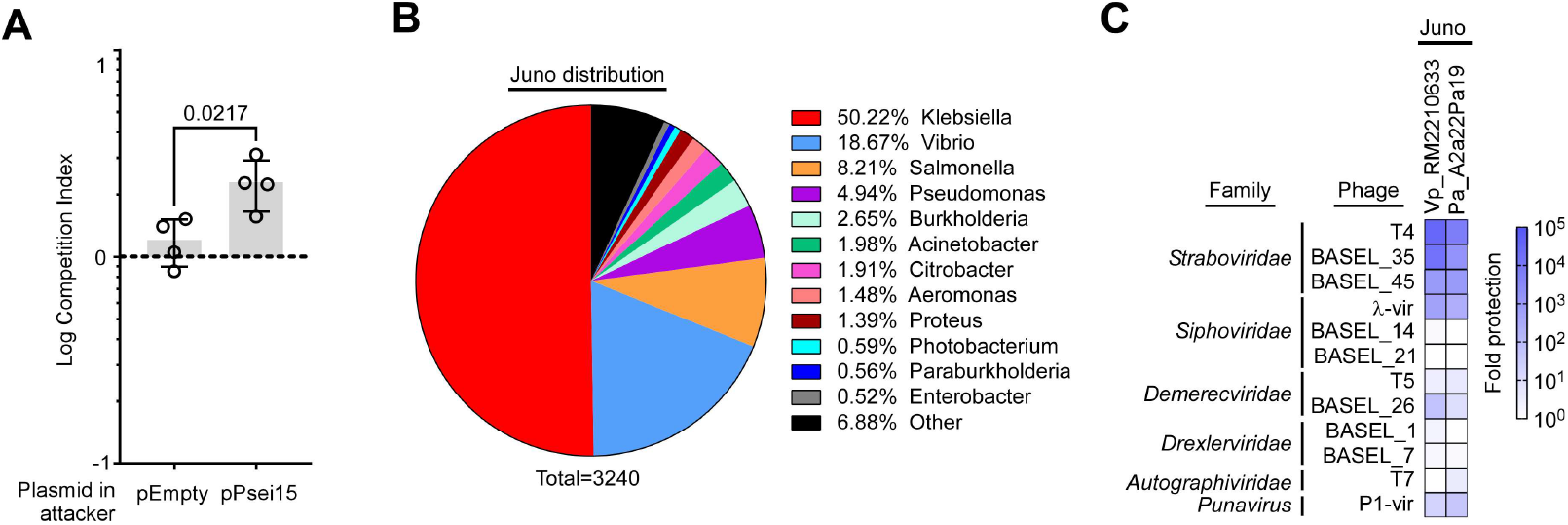
*P. agglomerans* variable genomic regions contain additional antibacterial T6SS effectors and anti-phage defense systems. **(A)** Competition between parental wild-type *Pab* prey strains and attacker *Pab* strains harboring the indicated plasmids, either empty or containing the predicted T6SS effector *pse15* together with its predicted promoter region and downstream immunity gene (pPsei15). Data are shown as Log competition index derived from the ratio of prey CFUs collected at 5 h after co-incubation with a T6SS^−^ (Δt*ssA*) attacker strain to those at the beginning of the experiment, divided by the same ratio for co-incubation with the wild-type attacker strains. The statistical significance between the samples was assessed using an unpaired, two-tailed Student’s t-test. Data are shown as the mean ± SD; n = the averages of 4 independent competition assays. **(B)** Pie chart showing the distribution of Juno homologs across bacterial genera. **(C)** The −Log EOP, indicating the reduction in plaque numbers (orders of magnitude) determined for *E. coli* expressing the indicated Juno homologs from an arabinose-inducible plasmid when challenged with the indicated coliphages, compared to *E. coli* containing an empty plasmid. The data shown are the average of 3 independent experiments.

### Using P. agglomerans hotspots to identify additional anti-phage defense systems

Next, because our findings indicate that *P. agglomerans* VRs contain a vast repertoire of anti-phage defense systems, we set out to determine whether investigating their cargoes could reveal additional anti-phage defense systems. In the VR_7 of *P. agglomerans* A2a22Pa19, which contains a Pse6 T6SS effector and two known anti-phage defense systems (Gabija and a restriction-modification system) (Dataset S4), we identified a gene predicted to encode a VPA1262 family protein (WP_397323501.1). The function of this protein family remains unknown; however, we have previously observed the founding member of this family in another mobile element, a GMT island in *Vibrio parahaemolyticus* RIMD2210633 (WP_005477100.1) (39). Since GMT islands are also known to be hubs of anti-phage defense systems (39), we hypothesized that members of the VPA1262 family function as anti-phage defense systems and renamed them Juno (after the queen of the Roman gods, protector of women and the state). Notably, Juno homologs are widely distributed among diverse Gram-negative bacterial genera (Fig. 7B).

To determine whether Juno protects bacteria against invading bacteriophages, we cloned it from both *P. agglomerans* A2a22Pa19 and *V. parahaemolyticus* RIMD2210633 into a low-copy-number arabinose-inducible expression plasmid. We then introduced these plasmids into *E. coli*, which is used as a surrogate host to examine sensitivity against coliphage challenge. We recently employed a similar strategy to identify anti-phage defense systems in GMT islands (39). *E. coli* strains were challenged with 12 coliphages representing different bacteriophage families (Table S1). When we compared the number of plaques each coliphage produced on a lawn of *E. coli* containing the Juno-encoding plasmids or an empty plasmid (i.e., the fold protection), we observed a dramatic reduction of > 100-fold in the number of visible plaques that developed in the presence of Juno with four coliphages: T4, BASEL_35, BASEL_45, and λ-vir (Fig. 7C). Therefore, we conclude that Juno is a widespread family of anti-phage defense systems. Collectively, our results further demonstrate the “genomic armory” nature of *P. agglomerans* VRs, and suggest that they contain additional defensive and offensive mechanisms to be discovered.

## DISCUSSION

To thrive in competitive ecological niches populated by diverse rivals and predators, bacteria must evolve or acquire versatile molecular arsenals that enhance their fitness. This adaptation is frequently driven by the integration of MGEs acquired via horizontal gene transfer (39, 72–74). Here, we demonstrate that *P. agglomerans*, a phytopathogen and opportunistic human pathogen, harbors dynamic genomic hotspots in which offensive and defensive payloads are either co-localized or alternate among strains. These diverse genetic cargoes include antibacterial T6SS apparatuses and effectors, prophages, anti-phage defense systems, and antibiotic resistance genes, collectively equipping bacteria to compete with bacterial rivals and withstand diverse biological threats.

In previous work, we characterized the combinatorial presence of offensive (i.e., antibacterial T6SS effectors) and defensive (i.e., anti-phage defense systems) arsenals within the cargo of widespread MGEs termed GMT islands (39). While the phenomenon of co-localizing antibacterial T6SS effectors and anti-phage defense systems was initially documented only in GMT islands of *Vibrionaceae*, recent studies have revealed genomic islands harboring both antibacterial and anti-phage repertoires in *Serratia* (63) and *Pseudomonas* (61) genomes. Together with our current findings demonstrating the co-localization of antibacterial T6SS effectors and anti-phage defense systems in *P. agglomerans* hotspots, these observations suggest that “genomic armories” represent a widespread feature of the bacterial accessory genome. Moreover, they imply that the co-occurrence of antibacterial and anti-phage arsenals is not restricted to particular MGEs, but rather occurs more broadly in genomic hotspots into which various MGEs can be integrated.

Collectively, these insights argue for broadening the traditional concept of “defense islands” (75, 76) beyond genomic regions solely dedicated to anti-phage mechanisms, toward a more inclusive concept of “genomic armories” that also integrate antibacterial weaponry. This functional plasticity, reflected in hotspots that alternate between offensive and defensive payloads, suggests that both strategies can contribute similarly to bacterial fitness. Alternatively, it may simply reflect a biological realization of the classic principle that the best defense is often a good offense.

Notably, our findings likely underestimate the true repertoire and diversity of the offensive and defensive arsenals in the cargoes of *P. agglomerans* hotspots. We demonstrate that the cargoes harbor previously unknown T6SS effectors (i.e., Pse15) and anti-phage defense systems (i.e., Juno), and we predict that additional examples can be uncovered by investigating genes of unknown function within these hotspots.

The diversity of effector arsenals associated with the antibacterial T6SS1 in *P. agglomerans* relies heavily on orphan effectors that appear to have been horizontally acquired and whose presence in the genome is not conserved across genomes, likely because of the dynamic nature of cargoes within genomic hotspots. A similar phenomenon of dynamic repertoires of orphan T6SS effectors was recently described in other bacteria, including *V. parahaemolyticus* (58) and *Pseudomonas aeruginosa* (42). However, the *P. agglomerans* T6SS1 effector repertoire substantially exceeds previously described arsenals associated with a single T6SS in both quantity and diversity.

Furthermore, we note that the current analysis was restricted to VRs harboring a T6SS effector within the *Pab* genome. However, we predict that the *P. agglomerans* pan-genome contains additional hotspots that serve as “genomic armories”, housing undiscovered T6SS effectors and anti-phage defense systems. These may be identified by analyzing the genomic synteny of T6SS effector homologs in multiple *P. agglomerans* genomes, following the workflow described in this work. Therefore, future studies should aim to systematically identify and characterize the dynamic cargoes of the mobile, accessory *P. agglomerans* pan-genome and to map its comprehensive weapon arsenal. Given that the *P. agglomerans* T6SS1 is conserved across many *Pantoea* and *Erwinia* species (36, 56), this analysis should provide broader insights into the effector landscapes and armory dynamics across members of these bacterial families. Moreover, the simple workflow described in this work, including the identification of antibacterial T6SS effector repertoires followed by their employment as “hooks” to reveal “genomic armories”, can be used to characterize the accessory armor landscape in other, diverse bacterial taxa.

In conclusion, by exploring the genomic neighborhoods of orphan T6SS effectors, we characterized a dynamic repertoire of genomic hotspots within the *P. agglomerans* pan-genome. These hotspots house diverse mobile armories, equipping bacteria with offensive and defensive arsenals that can enhance fitness during interbacterial warfare and phage predation. Crucially, the co-localization or alternating nature of antibacterial effectors and anti-phage defense systems within these armories suggests a functional blurring between aggression and protection. This paradigm implies that the antibacterial T6SS should be conceptualized not merely as an antagonistic, offensive system, but as an integral arm of the broader bacterial immune system.

## MATERIALS AND METHODS

### Strains and Media

For a complete list of strains used in this study, see Table S2. *Escherichia coli* strains were grown in 2xYT (1.6% [w/v] tryptone, 1% [w/v] yeast extract, 0.5% [w/v] NaCl), lysogeny broth (LB, 1% [w/v] tryptone, 0.5% [w/v] yeast extract, and 1% [w/v] NaCl), or on LB agar (1.5% [w/v]) agar) plates at 37°C. The media were supplemented with chloramphenicol (10 μg/mL), gentamicin (50 μg/mL), or kanamycin (50 μg/mL) to maintain plasmids when needed. L-arabinose (0.001%, 0.1%, or 0.2% [w/v], as indicated) was added to induce expression from the P*bad* promoter. *Pantoea agglomerans pv. betae* (*Pab*) and its derivatives were grown in 2xYT, LB broth, or on LB agar plates supplemented with rifampicin (50 μg/mL) at 28°C. Chloramphenicol (35 μg/mL), gentamicin (50 μg/mL), or kanamycin (50 μg/mL) was added to the media to maintain plasmids when needed.

### Plasmid construction

For a complete list of primers and plasmids used in this study, see Table S3 and Table S4, respectively. All cloning steps were performed using the Gibson Assembly method (77).

For the pPse7 and pPsi7 constructs, the coding sequences (CDS) of Pse7 (WP_064703290.1) and Psi7 (WP_128088276.1), respectively, were PCR-amplified from *Pab* genomic DNA. The amplicons were inserted into the multiple cloning site (MCS) of pBAD^K^/Myc-His or pBAD33.1 under an L-arabinose-inducible promoter, in-frame with a C-terminal Myc-His tag or a FLAG tag, respectively.

To express Pse15 and Psi15, the sequence encompassing the promoter region (261 bp upstream of *pse15*) and CDS of Pse15 (WP_062758233.1) and Psi15 (WP_122459110.1) from *P. agglomerans* strain NBBC-01 was synthesized by TWIST Bioscience with 20 bp flanks homologous to the intended MCS insertion site of the destination pBBR1MCS2 plasmid lacking a constitutive promoter (pBBR1MCS2_nopro). The sequence was introduced into pBBR1MCS2_nopro.

For expression of Juno variants, the CDS of WP_397323501.1 from *P. agglomerans* strain A2a22Pa19 with 20 bp flanks homologous to pBAD33.1 destination plasmid was synthesized by TWIST Bioscience. The CDS of VPA1262 (WP_231113970.1) was amplified from *V. parahaemolyticus* RIMD2210633 genomic DNA. The amplification products were inserted into the MCS of pBAD33.1 under an L-arabinose-inducible promoter.

Whole-plasmid sequence validation of the resulting constructs was performed by Plasmidsaurus using Oxford Nanopore Technology. The plasmids were introduced into *E. coli* via electroporation or into *Pab* via conjugation. Transformants and transconjugants were selected on LB agar plates supplemented with the appropriate antibiotics.

### Construction of deletion strains

For the deletion of *pse7* and *psi7* in *Pab*, a 730 bp sequence upstream of *pse7* and a 737 bp sequence downstream of *psi7* were cloned into pFOG, a Gent^R^R6Kγ suicide plasmid with dual negative selection mediated by I-SceI and SacB (78). The resulting plasmid, pFOG:psei7 was introduced into *E. coli* DH5α (λ-pir) by electroporation, and then into *Pab* via conjugation. Transconjugants were selected for the first homologous recombination event on LB agar plates supplemented with gentamicin (50 μg/mL) and rifampicin (100 μg/mL), and then counter-selected on LB agar plates containing 15% (w/v) sucrose and 0.5 μg/mL anhydrous tetracycline (AHT) for the second homologous recombination event. The deletion was confirmed by PCR.

### Comparative proteomics

#### Protein secretion

*Pab* wild-type (WT) and Δ*tssA* strains were grown overnight in LB media in triplicate. The following day, bacterial cultures were normalized to an OD_600_ of 0.5 in 2xYT media and incubated at 28°C with shaking (210 rpm). After 4 h, volumes equivalent to 10 OD_600_ units were collected from each sample and centrifuged at 4 °C to pellet the bacteria. The resulting supernatants were filtered through 0.22 µm filters, and proteins were precipitated using the deoxycholate-trichloroacetic acid (DOC-TCA) method (79). The precipitated proteins were washed twice with cold acetone and then shipped to the Smoler Proteomics Center at the Technion, Israel, for analysis.

#### Proteolysis

The protein pellets were dissolved in 8.5 M Urea, 400 mM ammonium bicarbonate, and 10 mM DTT. Protein concentrations were estimated using Bradford readings. The samples were reduced (60°C for 30 min), modified with 35.2 mM iodoacetamide in 100 mM ammonium bicarbonate (room temperature for 30 min in the dark), and digested in 1.5 M Urea, 66 mM ammonium bicarbonate with modified trypsin (Promega) overnight at 37°C in a 1:50 (M/M) enzyme-to-substrate ratio. An additional trypsinization step was performed for 4 hours in a 1:100 (M/M) enzyme-to-substrate ratio. An additional trypsinization step was performed under the same conditions.

#### Mass spectrometry analysis

The resulting peptides were analyzed by LC-MS/MS using an Exploris 480 mass spectrometer (Thermo) fitted with a capillary HPLC (EV-1000, Evosep One (480-Ex1). The peptides were loaded onto a 15 cm, ID 150 µm, 1.9-micron Performance column EV1137 (Evosep). The peptides were then eluted with the built-in Xcalibur 30 SPD (44 min) method. Mass spectrometry was performed in a positive mode using repetitively full MS scan (m/z 380–985, resolution 60, 000) followed by DIA scans (10 Da isolation windows with 1 m/z overlap, and resolution 30, 000).

#### Data analysis

The mass spectrometry data were analyzed using the DIA-NN software version 1.9.2 (80, 81) against *Pantoea agglomerans* section of the NCBInr database, with minimal peptide length set to 7, Maximum number of missed cleavages set to 1, Cysteine carbamidomethylation enabled as a fixed modification, and protein N-terminal acetylation enabled as a variable modification. Peptide- and protein-level false discovery rates (FDRs) were filtered to 1%. Statistical analysis of the identification and quantization results was performed using Perseus 1.6.7 software (82).

### Pse7 secretion assays

*Pab* wild-type and Δ*tssA* strains containing an empty plasmid or a plasmid for the L-arabinose-inducible expression of Pse7-myc were grown overnight in 2xYT media 28°C; the media were supplemented with kanamycin (50 μg/mL) to maintain the plasmid. The cultures were then washed with fresh media and resuspended to a final OD_600_ of 0.15 in 5 mL of 2xYT supplemented with kanamycin and 0.001% L-arabinose, to induce protein expression from the plasmid. Bacterial cultures were incubated with constant agitation (220 rpm) at 28°C for 5 h. For expression fractions (cells), cells equivalent to 1 OD_600_ unit were collected, and cell pellets were resuspended in 40 μL of 2x Tris-glycine SDS sample buffer (Novex, Life Sciences) supplemented with 5% (v/v) β-mercaptoethanol. For secretion fractions (media), volumes equivalent to 10 OD_600_ units were filtered through a 0.22 μm filter, and the proteins were precipitated using the deoxycholate and trichloroacetic acid method (79). Precipitated proteins were pelleted and washed twice with cold acetone, prior to re-suspension in 20 μL of 100 mM Tris-Cl (pH = 8.0), 20 μL of Tris-glycine SDS sample buffer supplemented with 5% β-mercaptoethanol, and 1 μL of 1 N NaOH to maintain a basic pH. The expression and secretion samples were resolved on SDS-polyacrylamide gel electrophoresis (SDS-PAGE), transferred onto nitrocellulose membranes using Trans-Blot Turbo Transfer (Bio-Rad), and immunoblotted with anti-Myc antibodies (Santa Cruz Biotechnology; 9E10, sc-40) at a 1:1000 dilution. Proteins were visualized using enhanced chemiluminescence (ECL). The membrane was then stained with 0.1% Ponceau S to confirm comparable loading of the samples. At least three independent experiments were performed.

### Competition assays

The indicated attacker and prey strains were grown overnight in LB and then normalized to OD_600_ = 0.5. Attacker and prey cultures were mixed at a 5:1 (attacker:prey) ratio, and the mixtures were spotted (25 μL) in triplicate on LB agar plates supplemented, when required for expression from a plasmid, with the indicated concentration of L-arabinose. The plates were incubated at 28°C for 5h. The colony-forming units (CFU) of the prey strain were determined by spotting tenfold dilutions of the mixtures from the 0 h timepoint and those harvested from competition plates at the 5 h timepoint on prey-selective plates. At least three independent experiments were performed.

Where used, the competition index was calculated by first dividing the average prey CFU at 5 h by the average prey CFU at 0 h during competition against a T6SS^−^ attacker across four biological replicates. The resulting ratio was then divided by the corresponding ratio obtained for competition against the wild-type (T6SS^+^) attacker.

### Plaque assays

Phages listed in Table S1 were propagated on *E. coli* K12 MG1655 ΔRM strains. To determine whether Juno homologs affect bacterial sensitivity to coliphages, *E. coli* K12 MG1655 ΔRM strains harboring the indicated pBAD33.1-based plasmids were grown overnight in LB supplemented with chloramphenicol and 0.2% (w/v) D-glucose (to repress expression from the P*bad* promoter) at 37°C. An empty pBAD33.1 plasmid was used as a negative control. Overnight cultures were washed twice with fresh media to remove any remaining glucose, and then 350 μL of each culture was mixed with 7 mL of 0.7% (w/v) molten agar supplemented with 0.2% (w/v) L-arabinose, 10 mM MgSO_4_, and 5 mM CaCl_2_. The mixtures were poured onto 1.5% (w/v) agar plates supplemented with chloramphenicol and 0.2% (w/v) L-arabinose, then left to dry for 1.5 hours. Tenfold serial dilutions of all the phages were prepared, and 7.5 μL of each dilution was spotted on the dried plates. The plates were incubated overnight at 37°C. The following day, the plaques were counted, and plaque-forming unit (p.f.u/mL) values were calculated. For dilution spots in which no individual plaques were visible, but a faint zone of lysis was observed, the dilution was considered as having ten plaques (83). At least three independent experiments were performed.

### Construction of the Phylogenetic tree of *P. agglomerans* genomes

A phylogenetic tree of 40 complete *P. agglomerans* genomes was constructed using M1CR0B1AL1Z3R 2.0 (84). The tree was based on the concatenated sequences of 1, 000 core proteins shared among the indicated strains. Phylogenetic relationships were inferred using the maximum-likelihood method. Bootstrap support values, calculated from 1, 000 replicates, are shown as percentages on the corresponding branches.

### Identifying Pse homologs

BLASTP was used to identify Pse homologs in 40 complete *P. agglomerans* genomes, as previously described (45). The protein accession numbers used as queries are provided in Dataset S3. Hits were filtered using an E-value threshold of 10^−6^, and a minimum alignment coverage of 70% of the query sequence length.

### Classification of T6SS gene clusters in *P. agglomerans* genomes

T6SS gene clusters were classified based on the phylogenetic distribution of the conserved T6SS core component TssM (Fig. S2 and Dataset S3). TssM homologs were identified using RPS-BLAST against COG3523, with an E-value threshold of 10^-9^. The resulting protein sequences were aligned using CLUSTAL-Omega v1.2.4 (85).

The evolutionary relationships among TssM homologs were inferred using the maximum-likelihood method under the WAG+G amino acid substitution model (86), which had the lowest Bayesian information criterion (BIC) score among 56 amino acid substitution models evaluated. Phylogenetic analyses were performed using MEGA X (87).

### Identification of Juno homologs

A position-specific scoring matrix (PSSM) for Juno was generated using PSI-BLAST. Five iterations against the RefSeq protein database were performed using the Juno protein WP_397323501.1 from *P. agglomerans* A2a22Pa19 as the query. In each iteration, up to 500 hits meeting an E-value threshold of 10^−6^ and a minimum query coverage of 70% were included.

A local database containing RefSeq bacterial nucleotide and protein sequences was constructed (last updated on February 23, 2025). Juno homologs were identified in the local protein database using RPS-BLAST with the generated PSSM. Hits were filtered using an E-value threshold of 10^−6^ and a minimum alignment coverage of 70%. The taxonomic distribution of the identified homologs was determined from the genus annotations of the corresponding RefSeq records.

### Variable genomic region identification and analyses

#### Genome download and ANI filtering

A total of 371 RefSeq assemblies of *P. agglomerans* genomes were downloaded from NCBI on May 26, 2026. Average nucleotide identity (ANI) relative to the *Pab* reference genome (GCF_001662025.2) was calculated using the OrthoANI algorithm (88), as implemented in pyOrthoANI (89). Assemblies with ≥ 95% were retained, resulting in a collection of 363 genomes for subsequent analyses (Dataset S2). A subset of 40 complete genomes was used to identify conserved regions flanking the effector loci, define the boundaries and anchors of variable genomic regions, and determine the distribution of effector homologs.

#### Sequence variability profiling

For each of 19 candidate orphan T6SS effector loci in the *Pab* reference genome, a 100-kb window extending 50 kb upstream and downstream from the effector start coordinate was extracted. Each window sequence was aligned against each of the 40 complete *P. agglomerans* genomes using BLASTN in megaBLAST mode (E-value threshold 10^-12^). High-scoring segment pairs (HSPs) were selected using a greedy, bitscore-based algorithm. HSPs were sorted by bitscore and iteratively retained if they matched the selected strand orientation and overlapped previously retained HSPs by less than 10%. To ensure conservation of the flanking sequences, contigs were retained only if the selected HSPs spanned both halves of the 100 kb window (positions 1-50, 000 and 50, 001-100, 000). When multiple contigs from the same genome met these criteria, the contig with the highest summed bitscore across the selected HSPs was retained. Per-base alignment occupation and nucleotide identity matrices were generated for each effector window, and the mean nucleotide identity across the genome collection was calculated at each position.

Next, all annotated genes, including noncoding genes, from the *Pab* reference genome were aligned against the subset of 40 complete genomes using BLASTN in megaBLAST mode (E-value threshold 10^-5^). For each hit, the Similarity Index (SI) was calculated as bitscore/(1.85 × query_length), where 1.85 represents the expected bitscore per base pair for a full-length 100%-identical alignment under the applied BLAST settings. Hits detected on different contigs of the same genome were grouped, and only the highest-bitscore hit per genome was retained.

Then, for each gene in a 100-kb effector window, the mean SI and percentage presence were calculated across the genomes represented in the corresponding identity matrix. Genes with a mean SI ≥ 0.8 and 100% presence were classified as conserved. The nearest conserved genes upstream and downstream of each effector locus were selected as anchors (Table 2).

#### Identification of variable genomic regions in P. agglomerans genomes

Anchor pairs were identified within the *P. agglomerans* 363 genomes by BLASTN searches using the anchor CDS/RNA sequences as queries (coverage ≥ 70%, E-value ≤ 10-^30^), requiring both anchors on the same contig in consistent orientation and order. Genomes containing multiple valid anchor pairs were manually reviewed, and one representative pair per genome was selected. This procedure yielded 17 variable genomic regions for downstream analyses (Table 2).

#### Cargo extraction, clustering, and functional annotation

For each variable genomic region and genome, genes fully contained between the two anchors, excluding the anchor genes themselves, were extracted as cargo. Cargo regions containing ≥ 11 consecutive ambiguous bases (Ns) were excluded from analysis of the corresponding variable region. Cargo nucleotide sequences were clustered independently for each variable genomic region using CD-HIT-EST v4.8.1 (-c 0.85, -s 0.85, -n 6, -G 1, -g 1, -r 1) (65). Proteins were annotated using DefenseFinder (67), PADLOC (68), RPS-BLAST (90) against NCBI CDD (91) and in-house PSSMs (RIX (6), PIX (43), WHIX (57), Tme (24), Juno), mobileOG (92), BLASTP against *Pab* T6SS effector and immunity seed sequences (E-value ≤ 10^-6^, ≥ 30% identity, and ≥ 70% subject coverage), BLASTP against the AMRFinderPlus AMRProt core dataset (93), retaining only canonical antimicrobial resistance (AMR) hits.

Pseudogenes were excluded from functional categorization. Each remaining cargo gene was assigned a single functional category using a fixed, priority-ordered rule set based on these annotations and a curated list of CDD PSSM IDs for T6SS effectors, T6SS structural components, toxin-antitoxin systems, antibacterial immunity proteins, and secreted toxins (Dataset S6).

### Prophage identification

Eight representative cargo sequences harboring predicted prophage regions were selected for detailed prophage characterization: VR_Pse6 (GCF_055709975.1), VR_Pse7 (GCF_003860325.1), VR_Pse9b+12 (GCF_010667985.1 and GCF_014841465.2), VR_Pse9d (GCF_049788445.1), VR_Pse11c (GCF_039672395.1), and VR_Pse13 (GCF_024799805.1 and GCF_001662025.2). Each cargo sequence was screened for integrated prophages using PHASTEST (69) through its URL API in “lite” annotation mode, in which bacterial genes are annotated against Swiss-Prot and phage regions are identified using the PHASTEST prophage database. Cargo sequences were submitted individually as single-sequence FASTA files. Predicted prophage regions were classified according to the PHASTEST completeness score as “intact” (score > 90), “questionable” (70–90), or “incomplete” (< 70), with all regions receiving the maximum score of 150.

## Supporting information

Supplementary Figures, Tables, Dataset captions, and References

## DATA AVAILABILITY

The scripts used to conduct the sequence analyses described in this work are publicly available and can be found at https://github.com/hiksenialeo/pab-genomic-islands/

The mass spectrometry data have been deposited in the ProteomeXchange Consortium via the PRIDE partner repository with the dataset identifier PXD076092. The data can be downloaded via https://ftp.pride.ebi.ac.uk/pride/data/archive/2026/07/PXD076092.

## ACKNOWLEDGEMENTS

We thank Dr. Alexander Harms (ETH Zurich) for generously sharing the BASEL phage collection. This research was supported by Research Grant Award No. US-5748-25 from BARD, The United States – Israel Binational Agricultural Research and Development Fund to DS; the European Research Council – Horizon Europe research and innovation programme (grant number 101169966) to DS; and by the Israel Science Foundation (ISF grant number 1362/21 to DS and EB, and 1067/25 to DS). UQ was supported by the European Research Council – Horizon 2020 research and innovation programme (grant number 818878). KL received financial support from the ADAMA Center for Novel Delivery Systems in Crop Protection, Tel Aviv University. CMF was supported by a Gray Scholarship for Post-Doctoral Fellows, Gray Faculty of Medical and Health Sciences Award, Tel Aviv University. We also thank the Smoler Proteomics Center at the Technion for performing and analyzing the mass spectrometry data. During the development of the analysis code, the authors used Cursor (Anysphere, Inc.), an AI-assisted coding environment powered by Anthropic’s Claude large language model, to help write, refactor, and debug scripts. All AI-assisted code was reviewed, tested, and validated by the authors, who take full responsibility for its correctness and for the work presented.

