## Supplementary Figures, Tables, Dataset captions, and References for "*Pantoea agglomerans* T6SS effectors reside in hotspots with combinatorial offensive and defensive arsenals"

**Supplementary Figures S1-S7**

**Supplementary Tables S1-S4**

**Supplementary Datasets S1-S6 (captions)**

**Supplementary References**

#### Supplementary Figures

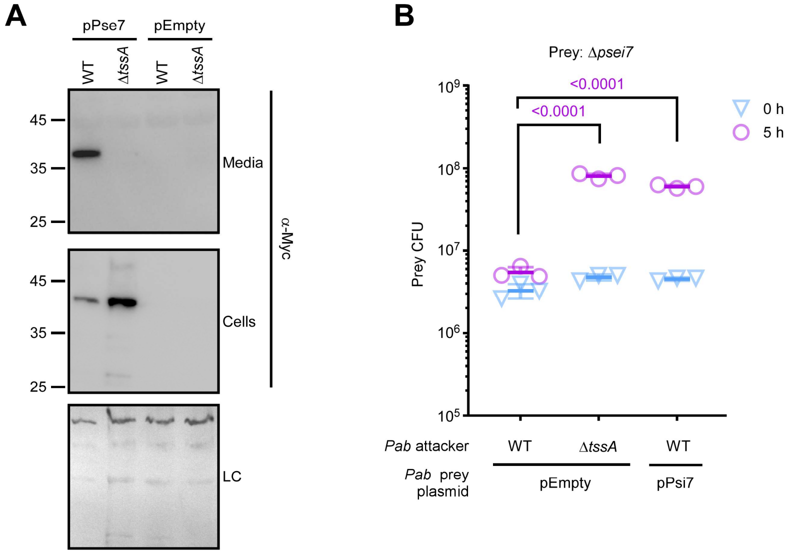


**Figure S1. Pse7 and Psi7 are an antibacterial effector and immunity pair. (A)** Expression (cells) and secretion (media) of C-terminal Myc-tagged Pse7 expressed from an arabinose-inducible plasmid (pPse7) in wild-type (WT) or Δ*tssA* (T6SS1$^{-}$) *Pab* strains grown for 5 h at 28°C in LB media supplemented with kanamycin and 0.001% L-arabinose. Total protein in the Cells fraction is shown as a loading control (LC). **(B)** Viability counts (CFU) of *Pab* prey strains in which the genes encoding Pse7 and Psi7 were deleted (Δ*psei7*), harboring an empty plasmid (pEmpty) or a plasmid for the arabinose-inducible expression of Psi7 (pPsi7), before (0 h) and after (5 h) co-incubation with the indicated *Pab* attacker strains on LB plates supplemented with 0.1% (wt/vol) L-arabinose at 28°C. The statistical significance between samples at the 5-h time point was assessed using one-way ANOVA followed by Dunnett's multiple comparisons test. Data are shown as the mean ± SD; n = 3. In panels A and B, results from representative experiments out of at least three independent experiments are shown.

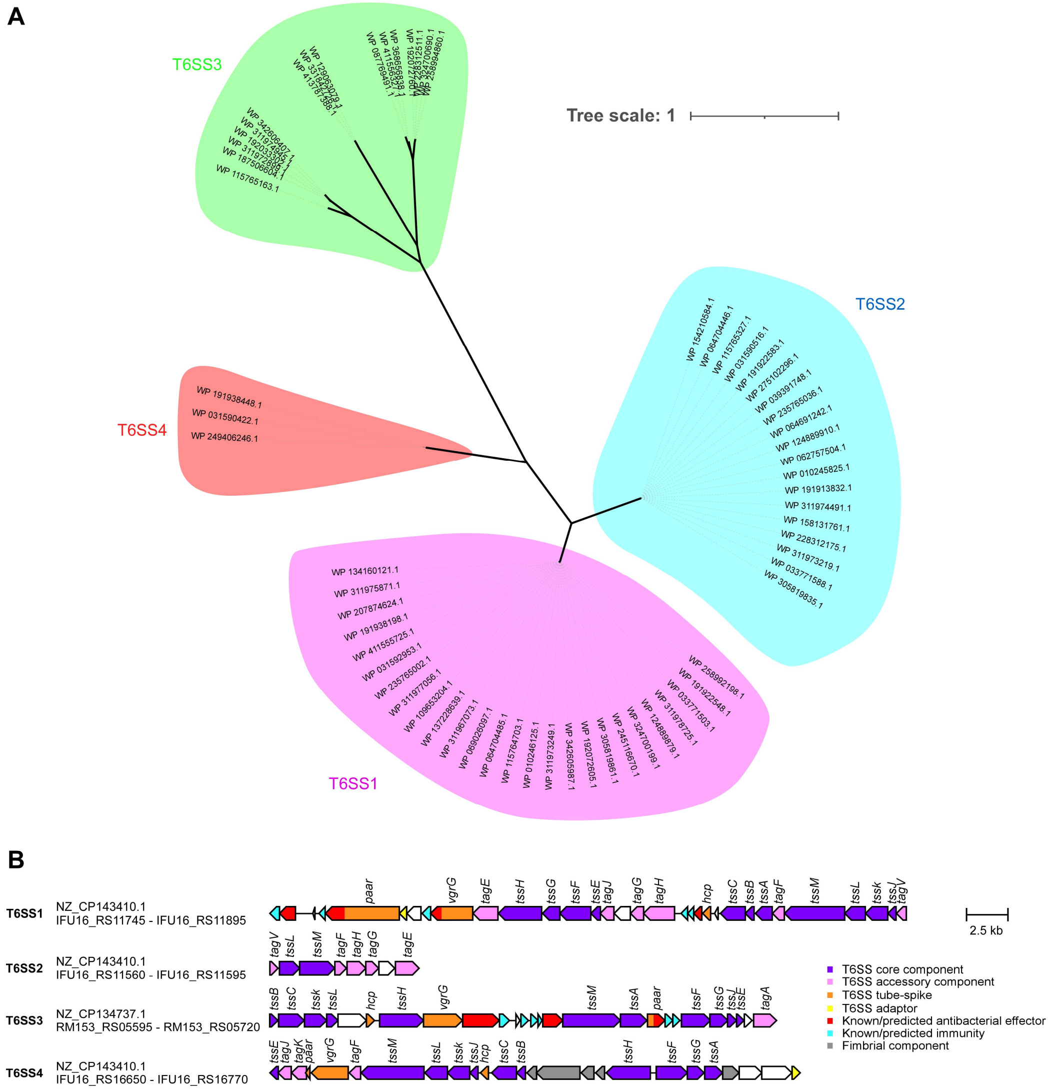


**Figure S2. The *P. agglomerans* pangenome contains four T6SS gene clusters. (A)** Phylogenetic distribution of the T6SS core component TssM encoded within 40 complete *P. agglomerans* genomes. The evolutionary history was inferred using the maximum-likelihood method and the Whelan and Goldman model. A discrete Gamma distribution was used to model evolutionary rate differences among sites. The phylogenetic tree was drawn to scale, with branch lengths measured in the number of substitutions per site. Evolutionary analyses were conducted in MEGA X. **(B)** Representative T6SS gene clusters found in *P. agglomerans* genomes. The GenBank accession numbers and locus tags of the cluster borders are provided.

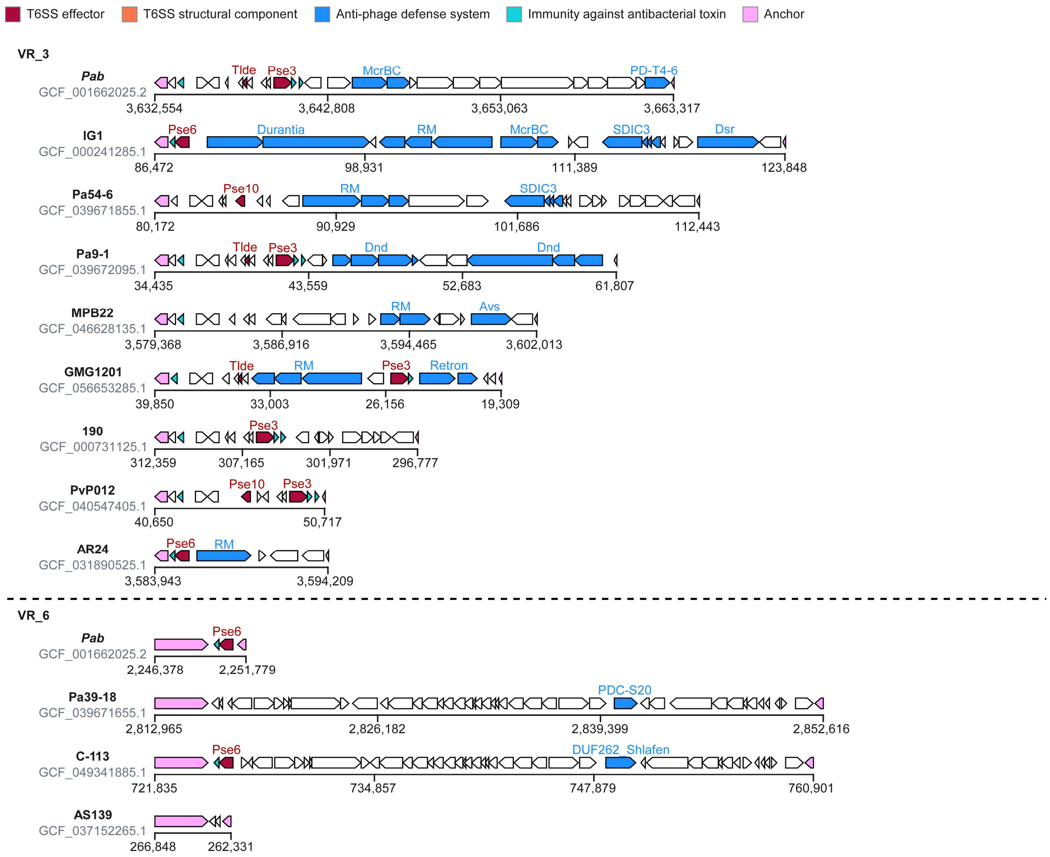


**Figure S3. Representative cargoes of the VR_3 and VR_6 variable genomic regions.** *P. agglomerans* strain names and NCBI RefSeq assembly numbers are denoted on the left.

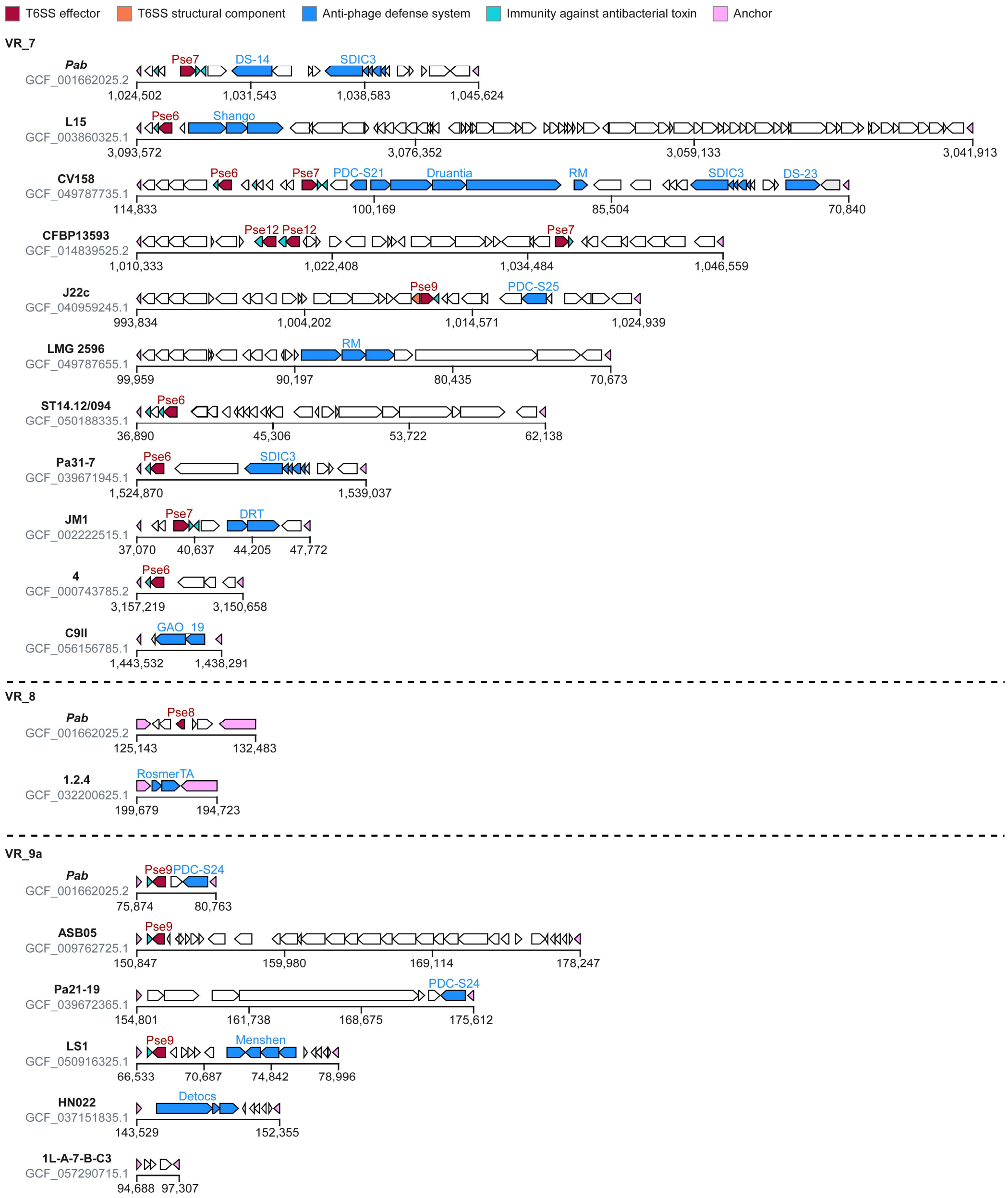


**Figure S4. Representative cargoes of the VR_7, VR_8, and VR_9a variable genomic regions.** *P. agglomerans* strain names and NCBI RefSeq assembly numbers are denoted on the left.

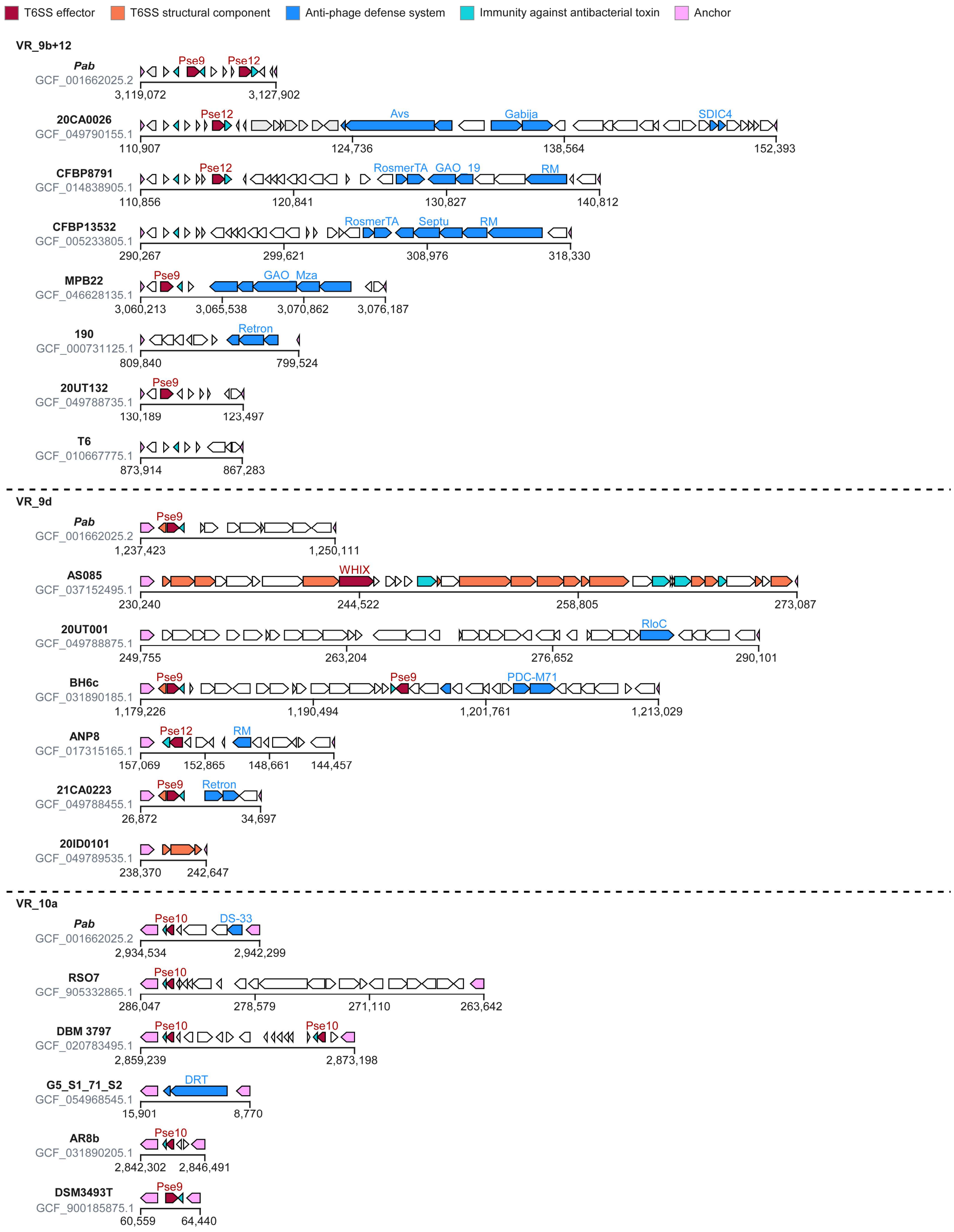


**Figure S5. Representative cargoes of the VR_9b+12, VR_9d, and VR_10a variable genomic regions.** *P. agglomerans* strain names and NCBI RefSeq assembly numbers are denoted on the left.

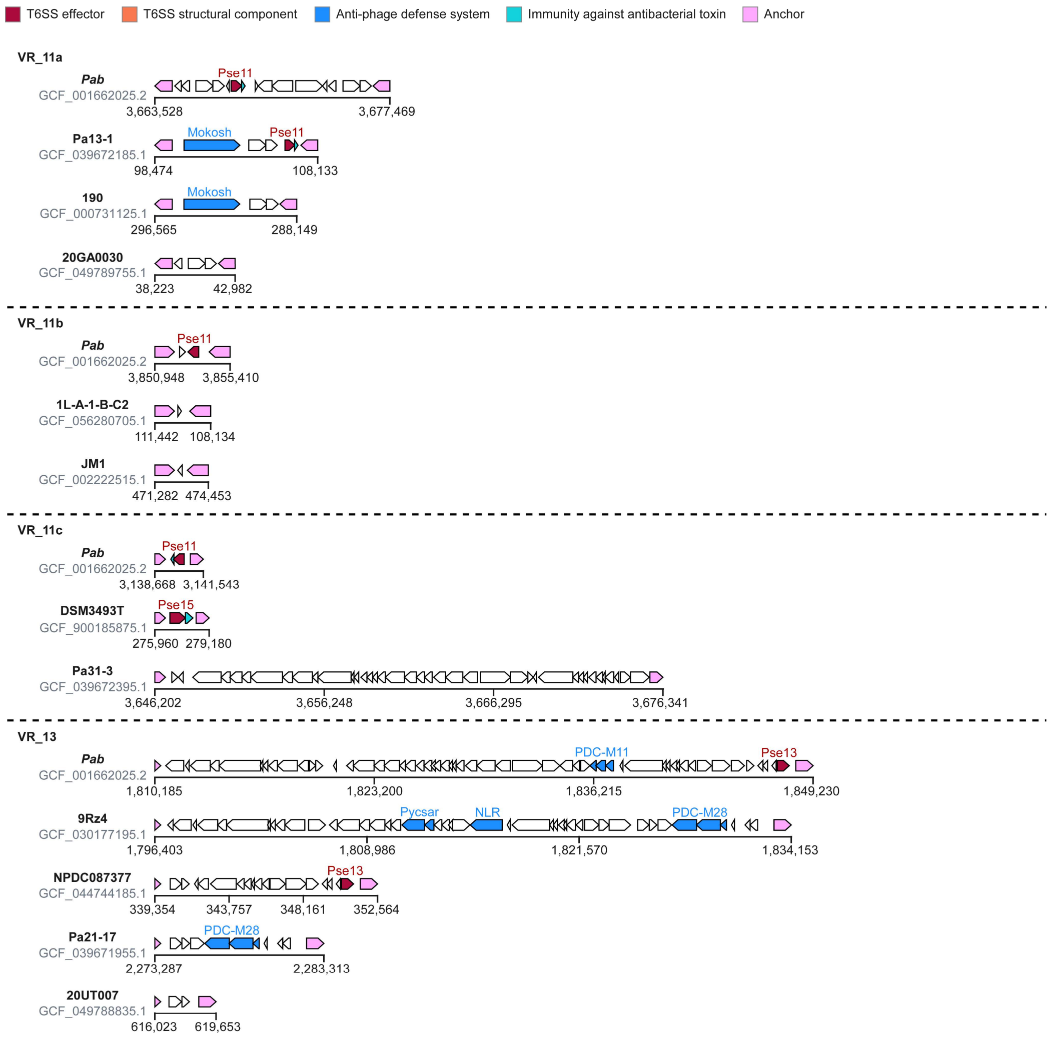


**Figure S6. Representative cargoes of the VR_11a, VR_11b, VR_11c, and VR_13 variable genomic regions.** *P. agglomerans* strain names and NCBI RefSeq assembly numbers are denoted on the left.

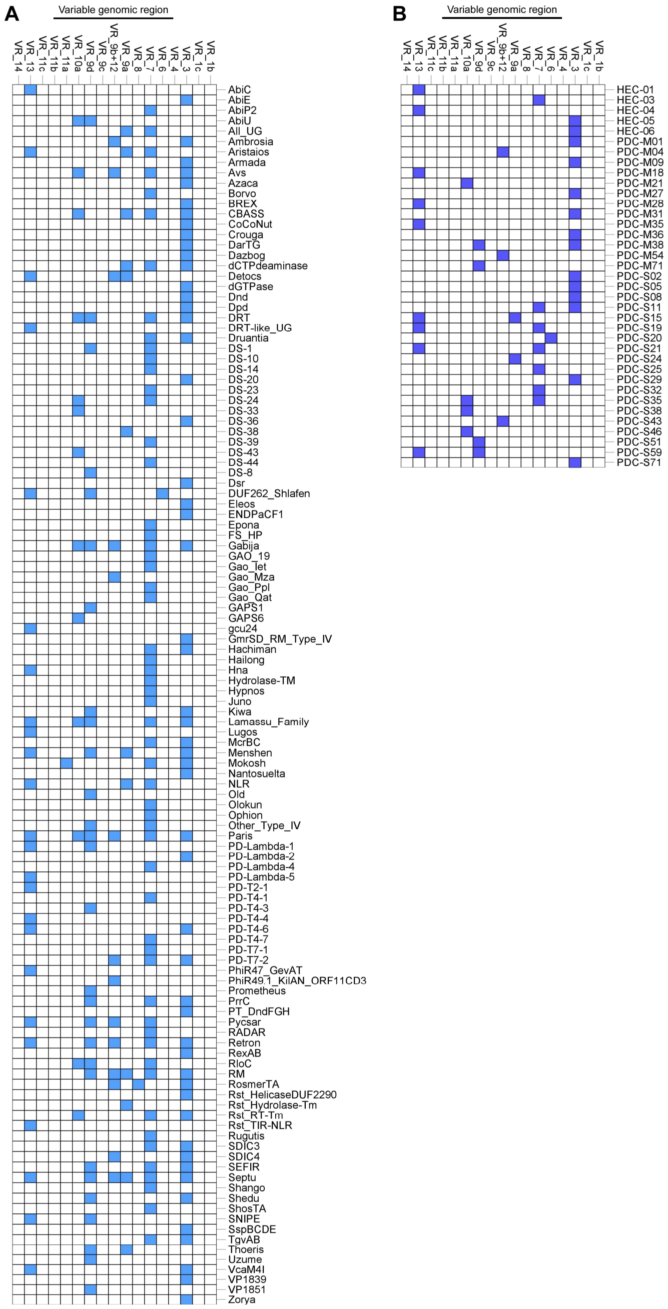

**Figure S7. Variable genomic regions in *P. agglomerans* are hubs for anti-phage defense systems.** Distribution of specific anti-phage defense systems, either verified **(A)** or predicted **(B)**, identified within cargoes of the indicated variable genomic regions. The analysis includes the anti-phage defense system Juno that was identified in this study, as detailed in the main text.

### Supplementary Tables

#### Table S1. List of bacteriophages used in this study.

| Phage | Family | Source |
|---|---|---|
| **T4** | *Straboviridae* | Lab collection |
| **T5** | *Demerecviridae* | Lab collection |
| **T7** | *Autographiviridae* | Lab collection |
| **P1-vir** | *Punavirus* | Lab collection |
| **λ-vir** | *Siphoviridae* | Lab collection |
| **Basel 01** | *Drexlerviridae* | (16) |
| **Basel 07** | *Drexlerviridae* | (16) |
| **Basel 14** | *Siphoviridae* | (16) |
| **Basel 21** | *Siphoviridae* | (16) |
| **Basel 26** | *Demerecviridae* | (16) |
| **Basel 35** | *Straboviridae* | (16) |
| **Basel 45** | *Straboviridae* | (16) |

**Table S2. List of bacterial strains used in this study.**

| Strain | Genotype | Use | Source |
|---|---|---|---|
| *Escherichia coli DH5α* | – φ80lacZΔ*M15* Δ(*lacZYA-argF*)*U169 recA1 endA1 hsdR17*(*rK– mK+*) *phoA supE44 λ– thi-1 gyrA96 relA1* Δ(*argF-lac*)169 pir+ | Used as a conjugation helper strain when harboring the plasmid pRK2073 | Lab stocks |
| *Escherichia coli DH5α (λ-pir)* | F– φ80lacZΔ*M15* Δ(*lacZYA-argF*)*U169 recA1 endA1 hsdR17*(*rK– mK+*) *phoA supE44 λ– thi-1 gyrA96 relA1* Δ(*argF-lac*)169 pir+ | Plasmid maintenance, cloning, protein expression | Obtained from Eric V. Stabb |
| *Escherichia coli* Neb5α | F- f80lacZDM15D(lacZYA-argF) U169 deoR recA1 endA1 hsdR17 (rk-, mk+) gal - phoA supE44 l- thi -1 gyrA96 relA1 | Plasmid maintenance and cloning | NEB |
| *Escherichia coli* K12 MG1655 ΔRM | Δmrr-hsdRMS-mcrBC ΔmcrA = ΔRM with pBR322_ΔPtet F(pifA::zeoR) | Bacteriophage propagation and plaque assays | (16) |
| *Pantoea agglomerans pv. betae 4188* (*Pab*) | Wild-type | Template for PCR amplification (gDNA), secretion assays, competition assays | Obtained from Isaac Barash |
| *Pab* Δ*tssA* | Deletion of A7P61_RS16010 | Secretion assays and competition assays | (1) |
| *Pab* Δ*psei7* | Deletion of *A7P61_RS09130* and *A7P61_RS09135* | Secretion assays and competition assays | This study |

##### Table S3. List of primers used in this study.

| Primer | Sequence (5'-> 3')' | Use |
|---|---|---|
| AH1F | TTAAGAAGGAGATATACATATGATTAACTATCTCAATACGTTGTTTGCAG | To amplify *juno* (*vpa1262*) from *Vibrio parahaemolyticus* RIMD2210633 for cloning into pBAD33.1 using Gibson Assembly |
| AH1R | ATCCGCCAAAACAGCCAAGCTTTTAGTGCGTGGTGAGTGTGTTC | |
| TM155R | CATATGTATATCTCCTTCTTAAAGTTAAAC | To amplify the pBAD33.1 vector backbone for Gibson Assembly to construct pJuno$^{Vp_RIMD2210633}$ |
| TM491F | AAGCTTGGCTGTTTTGGCGGATG | |
| T1639 | CCTGATACAGATTAAATCAGAACG | To amplify the pBAD33.1 vector backbone for Gibson Assembly to construct pJuno$^{Pa_A2a22Pa19}$ |
| T0339 | CATATGTATATCTCCTTCTTAAAGTTAAACAAAATTATTTCTAGAG | |
| T3399 | ACTGGGCTATCTGGACAAGGG | To amplify pFOG backbone for Gibson Assembly |
| T3400 | CAGCTTTTGTTCCCTTTAGTGAGGG | |
| T3401 | GCTACCTGCTTTCTCTTTGCGCTTGC | To amplify insert in pFOG MCS for colony screening |
| T3402 | TATGACCATGATTACGCCAAGCGCGC | |
| T3405 | CTAAAGGGAACAAAAGCTGATAGGTCATTAGAGGTGAACCTATATACCCC | To amplify a region upstream of *pse7* from the *Pab* genome for constructing pFOG:psei7 |
| T3406 | TCCGATATCATCTCTATCCTCAGTTATATGTTAAGCTGCCT | |
| T3407 | GAGGATAGAGATGATATCGGAAAATATTCATTAGCGACTTTATAAAACATGAGAGTG | To amplify a region downstream of *psi7* from the *Pab* genome for constructing pFOG:psei7 |
| T3408 | GTCCAGATAGCCCAGTCATGGCCTCGGCGAGCA | |
| T3380 | TAACAGGAGGAATTAACCATGGAAGTTATCCCTAGCGATATACAGTCA | To amplify *pse7* for constructing pPse7 in a pBAD$^{K}$/Myc-His backbone |
| T3383 | TTCGGGCCCAAGCTTTTGCTGAACTCCATACCAAACATTATACATCA | |
| T3534 | CGTCGTCATCCTTGTAATCTTTATTTAATCGAAGAAT | To amplify *psi7* for constructing pPsi7 in a pBAD33.1 backbone |
| T3535 | CTTTAAGAAGGAGATATACATATGATATCGGAAAATAAAGC | |
| T2234 | ATGCCATAGCATTTTTATCCATAAGATTAGCGG | To amplify insert in pBAD-based vector MCS for colony screening |
| T2235 | GATTTAATCTGTATCAGGCTGAAAATCTTCTCTCATC | |
| T4584 | TCACTGCCCGCTTTCCAGTC | To amplify pBBRMSC2 backbone without a promoter for Gibson Assembly |
| T4585 | ACTAGTTCTAGAGCGGCCG | |
| T4450 | GTAATCGGTACCCAGCTTTTGTTCCCTTTAG | To amplify the pBBRMSC2 backbone without a promoter for circularization through Gibson Assembly to construct an empty plasmid control, PBBR1MCS2_nopro |
| T4451 | GCTGGGTACCGATTACAAGGATGACGACG | |
| T3881 | ATGCATGCGCCCAATACG | To amplify the insert in pBBRMSC2 MCS for colony screening |
| T4192 | GGGTTTTCCCAGTCACGACGTTGTAAAACG | |

**Table S4. List of plasmids used in this study.**

| Plasmids | Origin/ antibiotic resistance | Description | Use | Source |
|---|---|---|---|---|
| pBAD33.1 | ori15a/CmR | L-arabinose-inducible expression vector | Used for cloning and arabinose- inducible expression | Addgene #36267 |
| pJuno$^{Vp_RIMD2210633}$ | ori15a/CmR | pBAD33.1 containing *vpa1262* from *Vibrio parahaemolyticus* RIMD2210633 in its MCS (WP_005477100.1) | Expression of Juno$^{Vp_RIMD2210633}$ in plaque assays | This study |
| pJuno$^{Pa_A2a22Pa19}$ | ori15a/CmR | pBAD33.1 containing Juno from *Pantoea agglomerans* strain A2a22Pa19 in its MCS (WP_397323501.1) | Expression of Juno$^{Pa_A2a22Pa19}$ in plaque assays | This study |
| pPsi7 | ori15a/CmR | pBAD33.1 containing *psi7* from *Pab* with a C-terminal FLAG tag (WP_128088276.1) | Expression of Psi7 in competition assays | This study |
| pBAD$^{K}$/Myc-His | pBR322/KanR | L-arabinose-inducible expression vector | Used for cloning and arabinose- inducible expression | (17) |
| pPse7 | pBR322/KanR | pBAD$^{K}$/Myc-His containing Pse7 from *Pab* with a C-terminal Myc-His tag (WP_064703290.1) | Expression of Pse7 in competition and secretion assays | This study |
| pBBR1MCS2_nopro | pBBR1/KanR | A derivative of pBBR1MCS2 vector with no constitutive promoter | Used for cloning and expression of genes under natural promoters | This study |
| pPsei15 | pBBR1/KanR | pBBR1MCS2_nopro vector containing *pse15* (WP_062758233.1) and *psi15* (WP_022626094.1) from *P. agglomerans* strain NBBC-01 with their natural promoter region (261 bp upstream of *pse15*) | Expression of Pse15 and Psi15 from their natural promoter in competition assays | This study |
| pFOG | RR6Kγ/GentR | A suicide vector with dual negative selection mediated by I-SceI and SacB | Used for deletion strain construction | Obtained from Dirk Bumann (18) |
| pFOG:psei7 | RR6Kγ/GentR | pFOG containing 730 bp sequence upstream of *pse7* and 737 bp sequence downstream of *psi7* in its MCS | Used for deleting psei7 in *Pab* | This study |
| pRK2073 | OriV/SpecR | A helper plasmid carrying conjugal transfer genes of RK2 | Used in the conjugation helper strain | Lab stocks |

#### Supplementary Datasets (captions)

**Dataset S1. Comparative proteomics analysis of the *Pab* T6SS1 secretome by mass spectrometry.**

**Dataset S2. OrthoANI analysis of *P. agglomerans* RefSeq genomes.**

**Dataset S3. Distribution of Pse homologs and T6SS gene clusters in complete *P. agglomerans* genomes.**

**Dataset S4. Analysis of cargo sequences identified within *P. agglomerans* variable genomic regions.**

**Dataset S5. PHASTEST analysis of prophage regions within representative variable genomic region cargo sequences.**

**Dataset S6. List of PSSMs used to categorize variable region cargo genes.**
